# Unravelling consensus genomic regions associated with quality traits in wheat (*Triticum aestivum* L.) using meta-analysis of quantitative trait loci

**DOI:** 10.1101/2021.11.24.469810

**Authors:** Santosh Gudi, Dinesh K Saini, Gurjeet Singh, Priyanka Halladakeri, Mohammad Shamshad, Mohammad Jafar Tanin, Pradeep Kumar, Achla Sharma

## Abstract

A meta-analysis of quantitative trait loci (QTLs) associated with following six major quality traits (i) arabinoxylan, (ii) dough rheology properties, (iii) nutritional traits, (iv) polyphenol content, (v) processing quality traits, and (vi) sedimentation volume was conducted in wheat. For this purpose, as many as 2458 QTLs were collected from the 50 mapping studies published during 2013-20. Of the total QTLs, 1126 QTLs were projected on to the consensus map saturated with 2,50,077 markers resulting into the identification of 110 meta-QTLs (MQTLs) with average confidence interval (CI) of 5.6 cM. These MQTLs had 18.84 times reduced CI compared to CI of initial QTLs. Fifty-one (51) MQTLs were also verified with the marker-trait associations (MTAs) detected in earlier genome-wide association studies (GWAS). Physical region occupied by a single MQTL ranged from 0.12 to 749.71 Mb with an average of 130.25 Mb. Candidate gene mining allowed the identification of 2533 unique gene models from the MQTL regions. *In-silico* expression analysis discovered 439 differentially expressed gene models with >2 transcripts per million (TPM) expression in grains and related tissues which also included 44 high-confidence candidate genes known to be involved in the various cellular and biochemical processes related to quality traits. Further, nine functionally characterized wheat genes associated with grain protein content, high molecular weight glutenin and starch synthase enzymes were also found to be co-localized with some of the MQTLs. In addition, synteny analysis between wheat and rice MQTL regions identified 23 wheat MQTLs syntenic to 16 rice MQTLs. Furthermore, 64 wheat orthologues of 30 known rice genes were detected in 44 MQTL regions. These genes encoded proteins mainly belonging to the following families: starch synthase, glycosyl transferase, aldehyde dehydrogenase, SWEET sugar transporter, alpha amylase, glycoside hydrolase, glycogen debranching enzyme, protein kinase, peptidase, legumain and seed storage protein enzyme.

**Main Conclusion:** Meta-QTL analysis in wheat for major quality traits identified 110 MQTLs with reduced confidence interval. Candidate gene mining and expression analysis discovered differentially expressed genes involve in quality traits.

## Introduction

Wheat (*Triticum aestivum* L.) is a globally grown cereal crop and is a major contributor of calories and protein to the human diet. Currently, wheat is widely consumed and processed into bread, noodles, cakes, pasta, beer, and other products. Wheat research has greatly contributed to its yield enhancement and disease resistance, but focus on quality of the produce took the back stage while enhancing yield. Hence, developing the high yielding varieties with enhanced quality characters is the foremost concern of the breeders (Nuttall et al. 2017). Improving end-use qualities is a tough endeavour because firstly, it is very difficult to measure the seed quality and rheological properties such as grain protein content (GPC), sedimentation rate (SDS), hectolitre weight (HW), 1000-grain weight (TGW), wet gluten content (WGC), dry gluten content (DGC), flour water absorption (FWA), dough development time (DDT), dough stability time (DST), mixing tolerance index (MTI), break down time (BDT) and kernel hardness (KH) as they are labour intensive and also require much seeds for analysis. Secondly, quality characters are complex traits governed by a plethora of gene networks that are largely influenced by several environmental conditions (Quraishi et al. 2017).

Quantitative trait locus (QTL) analysis has emerged as an effective approach for dissecting complex traits into component loci and studying the relative effects of the loci on the target trait (Doerge 2002). Since the first report on QTL analyses for wheat quality traits published in 1990s (Powell 1990), a plentiful of QTLs have been identified using different mapping populations to underpin the genetic architecture underlying end-use quality, including GPC (Huang et al. 2006; Mann et al. 2009), dough rheological properties (Huang et al. 2006; Mann et al. 2009), SDS (McCartney et al. 2006), falling number (FN) (Fofana et al. 2009) and starch pasting properties (Mohler et al. 2014). However, the rationality of these QTL mapping results is strongly influenced by the experimental conditions, type and size of mapping population, density of genetic markers, statistical methods used among others (Swamy et al. 2011). Thus, the practical implication of these QTLs for quality improvement via molecular QTL cloning and marker-assisted selection has been rather limited (Quraishi et al. 2017). Considering this challenge, it is desirable to identify QTLs that show major effect on target phenotype and are consistently detected across the multiple genetic backgrounds and environments.

Meta-analysis of available QTLs enable the identification of consensus and robust QTLs or MQTL regions that are most frequently associated with trait variation in diverse studies and reduce their confidence intervals (CIs) (Goffinet and Gerber 2000; Veyrieras et al. 2007). Software packages, such as Meta-QTL and BioMercator, facilitate meta-analysis of QTLs derived from independent studies by formulating and embedding specific sets of algorithms for exact evaluation and recalculation of the genetic position for the given QTLs (Wang et al. 2014). Among them, BioMercator is the most advanced and commonly used software for projection of initial QTLs on consensus map and identification of MQTLs. In wheat, meta-QTL analysis has already been conducted for different traits, including the ear emergence (S et al. 2009), fusarium head blight (Liu et al. 2009; Venske et al. 2019; Zheng et al. 2021), tan spot resistance (Liu et al. 2020), pre-harvest sprouting tolerance (Tyagi and Gupta 2012), abiotic (drought and heat) stress tolerance (Acuña-Galindo et al. 2015; Darzi-Ramandi et al. 2017; Kumar et al. 2020), yield (Yang et al.; Tyagi et al. 2015; Quraishi et al. 2017; Kumar Saini et al. 2021) and baking-quality and GPC (Quraishi et al. 2017). Quraishi et al. (2017) identified the six and eight MQTLs for GPC and baking quality traits, respectively, using the 155 initial QTLs collected from eight interval mapping studies. S**Error! Bookmark not defined.**ince, large number of QTLs have been reported after the meta-QTL analysis for quality traits by Quraishi et al. (2017) in wheat, the present study involving meta-QTL analysis was performed (based on interval mapping studies published during 2013-20) to supplement the list of MQTLs and candidate genes reported in the earlier MQTL study for quality traits (Quraishi et al. 2017). Further, the results of the meta-analysis were integrated with GWAS and transcriptomics studies to identify the promising genomic regions and important CGs, which affect quality traits in wheat. Further, by utilizing the synteny and collinearity of wheat with other cereals (Sorrells et al. 2003), ortho-MQTL analysis was conducted to see if the generated information can be transferred to other cereals such as rice. The findings of the present study may help in identification of diagnostic markers and aid marker-assisted breeding to improve quality traits in wheat.

## Material and methods

### Bibliographic search and QTL data cumulation

A comprehensive bibliographic search and data retrieval on the published papers from 2013 to 2020 enabled us to compile information from the available studies. From each independent study, the following information was obtained: (i) QTL name (wherever available), (ii) flanking or closely linked markers, (iii) peak position and CI, (iv) LOD score, (v) phenotypic variation explained (PVE) or R^2^ value for each QTL and (vi) type and size of the mapping population (S1 and S2 Tables). Wherever, information about the peak position of the QTL was missing, it was calculated as the mid-point of the two markers flanking the QTL, and when LOD scores of individual QTLs were unavailable from a particular study, LOD score of 3 was considered for all the QTLs identified in the concerned study.

All the quality traits were grouped into the following six major trait categories: (i) *Arabinoxylan* (*Ax*) recorded as total Ax, sucrose solvent retention capacity (SRC), sucrose softness equivalent SRC, water-extractable and water-unextractable Ax; (ii) *Dough rheology properties (DRP)* recorded as mixolab, mixograph, mixogram, alveograph, farinograph, water SRC, alkaline SRC, dough mixing, and pasting properties; (iii) *Nutritional traits (NT)* recorded as lipoxygenase activity, yellow pigment, Fe, Zn, β-glucan, protein, and starch content; (iv) *Polyphenol content* (*PPO*) recorded as polyphenol oxidase activity, flour colour, and loaf colour; (v) *Processing quality traits (PQT)* recorded as milling and baking qualities; and (vi) *Sedimentation volume* (*SV*) recorded as SV, sedimentation rate and zeleny SV (S3 Table).

### Construction of consensus map

An R package LPmerge (Endelman and Plomion 2014) was used to construct a consensus map (named as Wheat_Reference_GeneticMap-2021) by merging the following linkage maps: “Wheat_Composite_2004” (http://wheat.pw.usda.gov), “Wheat_Consensus_SSR_2004” (Somers et al. 2004); “Durum wheat integrated map” (Marone et al. 2013); and four SNP array based maps developed using “Illumina 9K iSelect Beadchip Array” (Cavanagh et al. 2013), “Illumina iSelect 90K SNP Array” (Wang et al. 2014), “Wheat 55K SNP array” (Winfield et al. 2016) and “AxiomR, Wheat 660K SNP array” (Cui et al. 2017). Markers flanking the initial QTLs identified from different mapping studies were also integrated on to the consensus map (S4 Table).

### QTL Projection, meta-QTL and ortho-MQTL analysis

For the QTLs with no CI available, CI (95%) was calculated by using the following population-specific equations (Darvasi and Soller 1997; Venske et al. 2019).

For RIL,

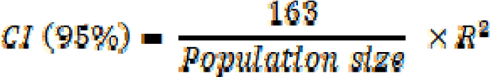

For DH,

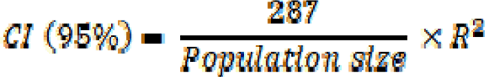

Further, careful examination and evaluation of individual QTLs for genetic map or QTL-related information led us to exclude 472 QTLs from the further analysis. Information on LOD score, PVE, genetic position (i.e., CI and peak position) from the remaining 1986 QTLs were compiled and used for projection on consensus map using the QTLProj tool of BioMercator V4.2. Meta-analysis was then performed for the individual chromosomes using Veyrieras two-step approach available in the software. In the first step, the best QTL model (used to ascertain the number of MQTL per chromosome) were computed by using the following five criteria: (i) Akaike Information Criterion (AIC), (ii) corrected Akaike Information Criterion (AICc), (iii) Akaike Information Criterion 3 (AIC3), (iv) Bayesian Information Criterion (BIC) and (v) Approximate Weight of Evidence Criterion (AWE). QTL model which had the least criterion value and zero delta value was considered as the best model for the analysis. In the second step, selected model was considered to determine the number of MQTLs on each chromosome (based on number of input QTLs on common genetic map), their consensus positions (based on variance of input QTL positions) and 95% CI (based on variance of input QTL intervals) (Sosnowski et al. 2012).

To investigate the ortho-MQTLs for quality characters, a previous meta-QTL study (Youlin et al. 2021) conducted in rice for starch pasting properties was used. To mine the ortho-MQTLs between wheat and rice, orthologous regions between the two species were investigated by using EnsemblPlants database. Candidate genes available in selected MQTL regions were used to search the corresponding rice genes. MQTL regions with at least 4 conserved genes present in wheat and rice genome were considered as the ortho-MQTL (Youlin et al. 2021).

### Delineating physical location and candidate gene mining in MQTL regions

Nucleotide sequences of the markers flanking the MQTLs were retrieved from either of the following databases/websites: (i) the GrainGenes (https://wheat.pw.usda.gov/GG3), (ii) Diversity array technology (https://www.diversityarrays.com) and (iii) CerealsDB (https://www.cerealsdb.uk.net/cerealgenomics/CerealsDB/indexNEW.php). These sequences were used to obtain the physical positions of markers by BLASTN searches against Wheat Chinese Spring IWGSC RefSeq v1.0 genome assembly available in EnsemblPlants database (http://plants.ensembl.org/index.html). Physical positions of some of the SNP markers were directly obtained from the JBrowse-WHEAT URGI database (https://urgi.versailles.inra.fr/jbrowseiwgsc/).

Peak physical positions of MQTLs were calculated by using the following formula:

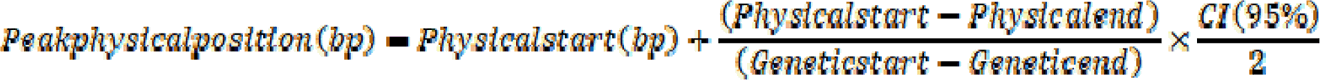

Gene models available in 2 Mb genomic region i.e., 1 Mb region on either side of the MQTL peak position were retrieved using the ‘BioMart’ tool of EnsemblPlant database (https://plants.ensembl.org/biomart/). Functional characterization and gene ontology (GO) analysis for available gene models was also carried out using the same tool BioMart.

### Expression analysis for gene models

An adaptable platform to create a gene expression interface **‘**Wheat expression browser’ powered by expVIP was used to analyse the expression of gene models available in the MQTL regions (Borrill et al. 2016). Following relevant gene expression datasets: (i) grain tissue-specific developmental time course (CS grain, grain dissection, early grain) (Gillies et al. 2012; Li et al. 2013; Pfeifer et al. 2014); (ii) grain tissue-specific expression at 12 days post anthesis (Pearce et al. 2015) and (iii) grain developmental time course with 4A dormancy QTL (Barrero et al. 2015) were used for this purpose. These studies provided the expression data on “morphological stages of developing wheat grain”, “inner pericarp, outer pericarp and endosperm layers from developing grain of bread wheat at 12 days post-anthesis”, “aleurone and starchy endosperm tissues of the wheat seed at aleurone layer development time of 6, 9 and 14-days post anthesis” and “candidate genes underlying the grain dormancy in wheat”, respectively.

Only the gene models with ≥ 2 TPM (transcripts per million) expression were considered in the present study (GP et al. 2013). Further, protein sequences of the known genes associated with quality traits (collected from the literature) were retrieved and BLASTP searches were made against the wheat genome database available in EnsemblPlants to find their physical positions. Then, the physical positions of these genes were compared with the physical coordinates of the MQTLs to determine their co-localization, if any.

### Validating MQTLs with MTAs identified in earlier GWA studies

Information from the 11 independent GWA studies on most stable and significant MTAs were collected to validate the MQTLs identified in the present study. These GWA studies involved one durum wheat population (with population size 194), four spring wheat populations (with population size ranging from 189 to 2038), three winter wheat populations (with population size ranging from 267 to 1325) and three mixed population of spring and winter wheat (with population size ranging from 163 to 4095). P**Error! Bookmark not defined.**hysical positions of each significant and stable SNPs associated with the trait were obtained from the respective studies and/or JBrowse-WHEAT URGI database (https://urgi.versailles.inra.fr/jbrowseiwgsc/). Finally, physical positions of these MTAs were compared with the physical coordinates of the MQTLs, to find any MQTL co-localizing with at least one MTA and such MQTLs were considered as a GWAS-validated/verified MQTL.

### MQTL characterization using cloned genes from the rice

Functionally validated rice genes associated with quality traits (involving starch and protein metabolism, embryo and endosperm development, sugar transportation, grain development, etc.) were collected from the available literature and their protein sequences were retrieved from the Rice Annotation Project Database (rap-db) (https://rapdb.dna.affrc.go.jp/index.html). The protein sequences of rice genes were then BLAST against wheat genome to identify their corresponding wheat homologues available in the MQTL regions.

## Results

### Characterization of QTL studies involving quality traits

Characteristics of 50 independent mapping studies involving 63 bi-parental populations were rigorously reviewed to compile information on available QTLs. In total, 2458 QTLs associated with quality traits were collected and they were found to be distributed unequally on all the wheat chromosomes (Fig 1a). Maximum number of QTLs were present on the sub-genome B (995 QTLs), while the minimum number of QTLs were present on the sub-genome D (607 QTLs). There were 45 sets of RIL populations (population size ranging from 83 to 171) and 18 sets of DH populations (population size ranging from 94 to 192) (Fig 1b, c; S1 and S2 Tables). The number of studies involving different DH and RIL populations, molecular markers and mapping methods are presented in Fig 1c, d and respectively. Most of the mapping studies used SSR as the markers and composite interval mapping (CIM) as method for QTL mapping, respectively (Fig 1d, e). Number of QTLs associated with the different quality traits also varied (from 36 QTLs for Ax content to 962 QTLs for DRP). PVE values and LOD score for individual QTLs ranged from 0.01 to 95.24 per cent with a mean of 9.31 per cent and from 0.99 to 55.32 with an average of 5.74, respectively (Fig 1f, g; S1 Table). Peak positions of the initial QTLs also varied (ranging from 0 to 678.2 cM with a mean of 108.18 cM) (Fig 1h).

**Fig 1.**
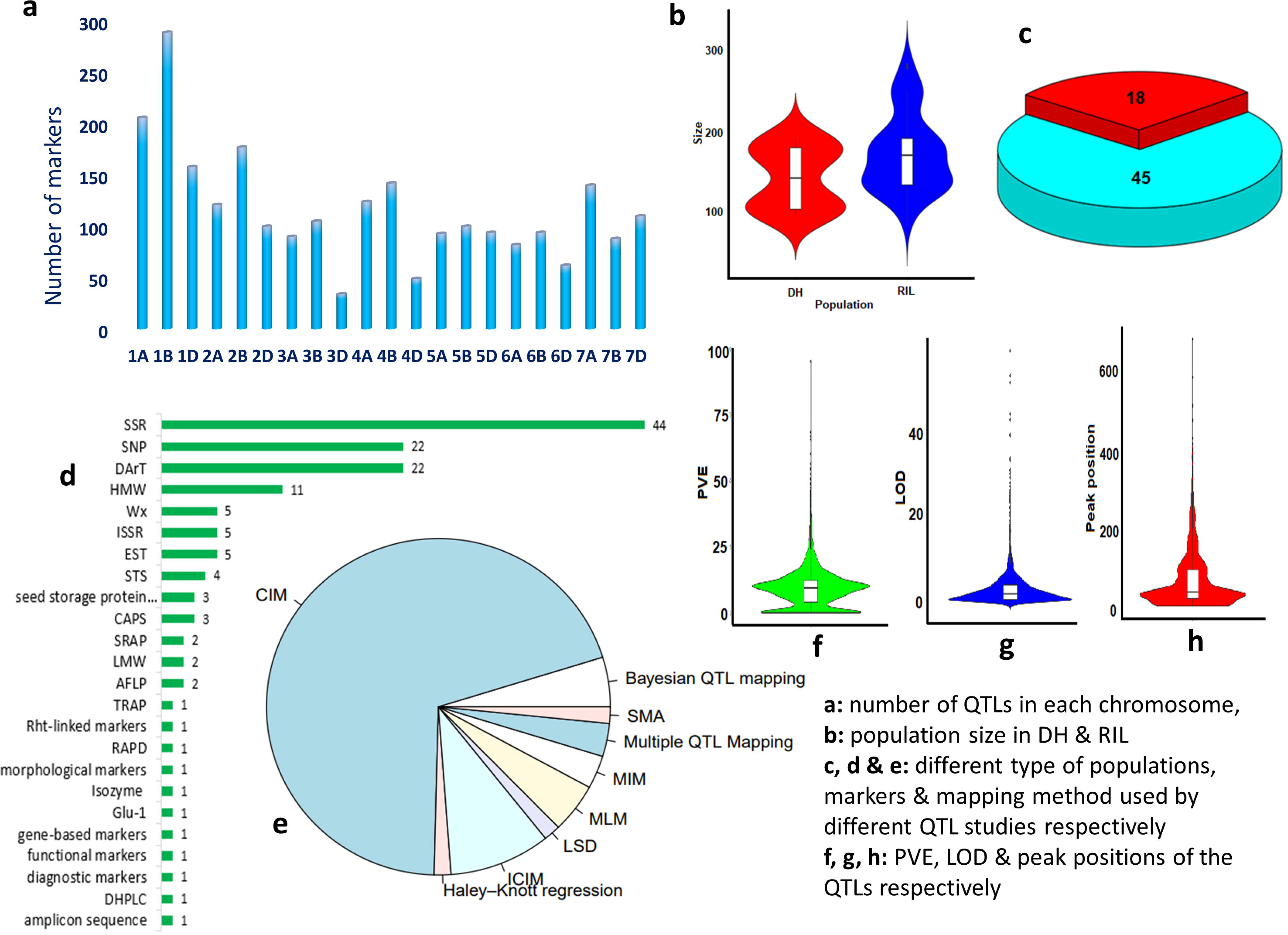
Salient features of the initial QTLs. (a) number of QTLs available from different chromosomes, (b) sizes of the populations utilized for mapping, (c) number of different types of populations, (d) different types of markers utilized for mapping, (e) methods employed for mapping, (f) PVE, (g) LOD, and (h) peak positions of the QTLs.

### Construction of consensus genetic map

The consensus map, “Wheat_Reference_GeneticMap-2021” showed significant variation for the genetic lengths of the individual linkage groups/chromosomes (ranging from 361.8 cM for 1A to 808.6 cM for 2A, with a mean of 649.4 cM) and for the number of markers positioned on a single chromosome (ranging from 2,225 on 4D to 21,253 on 3B with an average of 11908.4 markers per chromosome) (Fig 2). The total length of the consensus map measured 13,637.4 cM which included a total of 2,50,077 markers of different types such as AFLP, RFLP, SSR, SNP, etc. Since, different genetic maps with varying number and type of markers were used to construct the consensus map, the distribution of markers on the chromosomes was uneven and density of the markers was comparatively higher at the fore-end of the chromosome (Fig 2; S4 Table). Marker density on individual chromosomes ranged from 5.7 markers/cM on 4D to 46.2 markers/cM on 1A with a mean of 18.3 markers/cM on whole genome.

**Fig 2.**
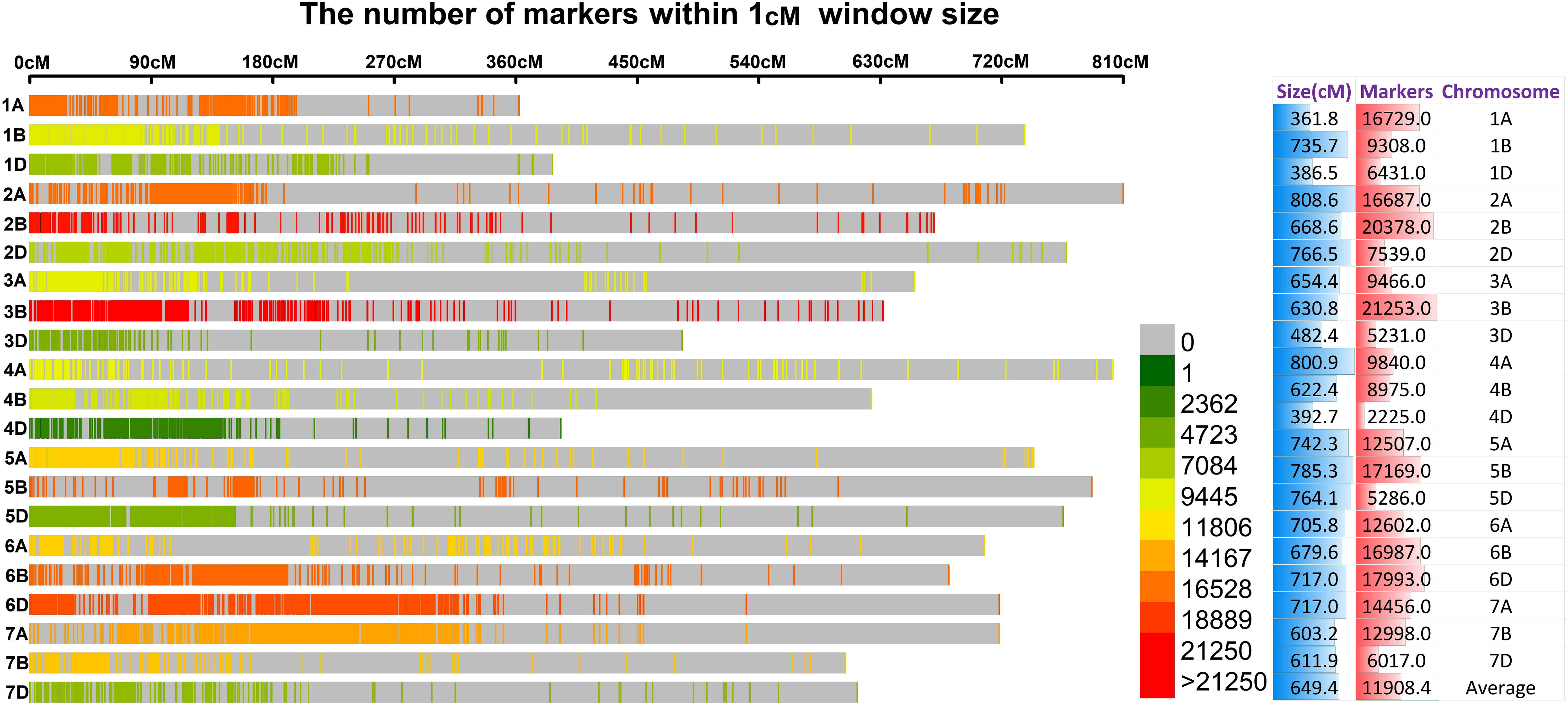
Consensus map, “Wheat_Reference_GeneticMap-2021” showing the marker density (number of markers/cM), chromosome length (cM) and number of markers in each chromosome.

### MQTLs identified for quality traits

Of the total 1986 QTLs with complete information available for analysis, only 1128 (56.79% QTLs were projected onto the consensus map. Among the 1128 QTLs, 1121 QTLs were grouped into 110 MQTLs, 7 QTLs remained as singletons (Table 1; Fig 3 and 4). Eight hundred and fifty-eight (850) QTLs were not projected onto the consensus map owing to either of the following reasons: (i) there were no common markers between the consensus and original maps, and (ii) the QTLs had low R^2^ values, resulting in a large CI. The number of MQTLs on individual chromosomes ranged from 2 (on chromosomes 5B, 6B and 7A) to 9 (on chromosome 1A) (Fig 3a; S5 Table). The number of clustered QTLs per MQTL ranged from 2 (in several MQTLs) to 85 (in MQTL1B.1) with 11 MQTLs involving at least 20 initial QTLs (Fig 3d; S5 Table). Of these 11 MQTLs, four were located on chromosome 1B and remaining each of the seven MQTLs were located on different chromosomes. Since, these 11 MQTLs integrated the QTLs from different populations, they were considered to be more stable and reliable MQTLs for wheat quality breeding. The chromosome-wise average CI (95%) for the initial QTLs and MQTLs (presented in the Fig 3c) ranged from 14.87 cM (1D) to 95.55 cM (3D) for QTLs and from 0.55 cM (1D) to 34.30 cM (2D) for MQTLs. The average CI of MQTLs (5.56 cM) was 18.84-fold less than that of initial QTLs (40.35 cM), and there were significant differences among different chromosomes (Fig 3c). The average CI of MQTLs reduced maximum for chromosome 5B (72.55 times) and minimum for chromosome 2D (1.99 times) compared to the CI of initials QTLs located on corresponding chromosomes.

**Fig 3.**
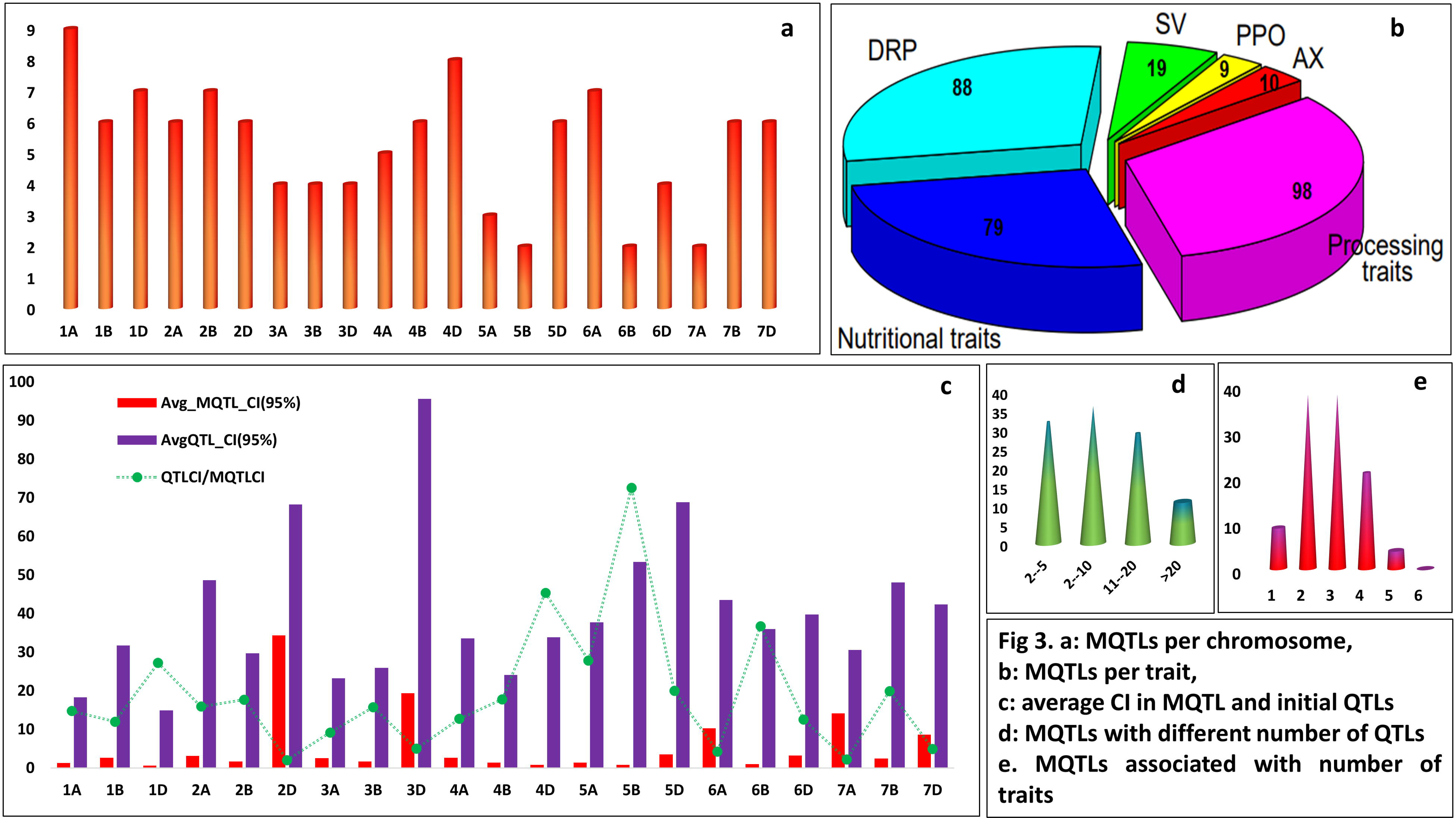
Characteristic features of the MQTLs. (a) distribution of MQTLs identified on different wheat chromosomes, (b) trait-wise distribution of the MQTLs, (c) average CIs of initial QTLs and MQTLs and fold reduction in CI values after meta-QTL analysis, (d) different number of QTLs involved in MQTLs, (e) MQTLs associated with different number of traits.

**Fig 4.**
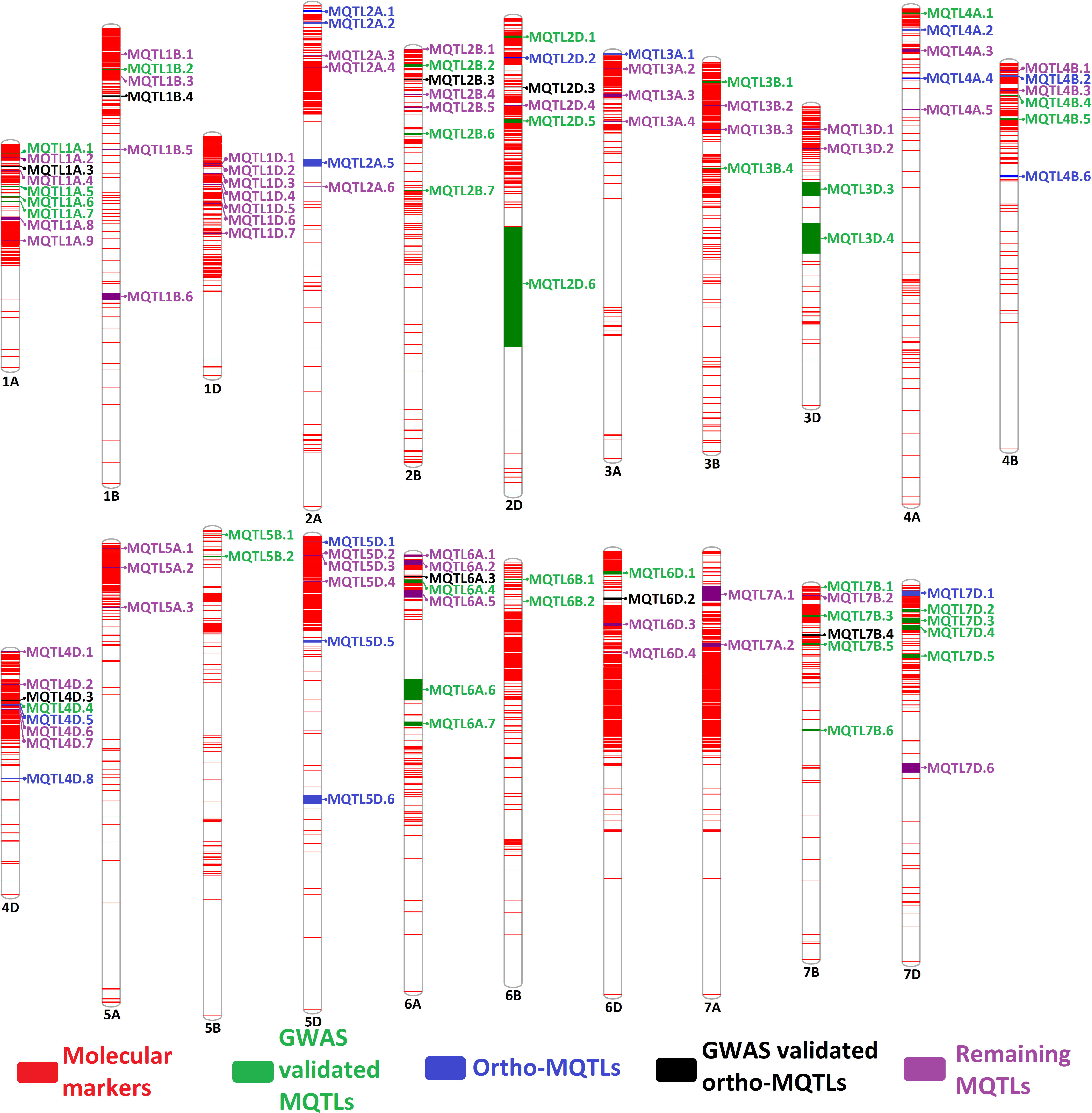
Distribution of different MQTLs on wheat chromosomes. Explanation for different colours utilized to represent the MQTLs are given at the bottom of the figure.

**Table 1.**
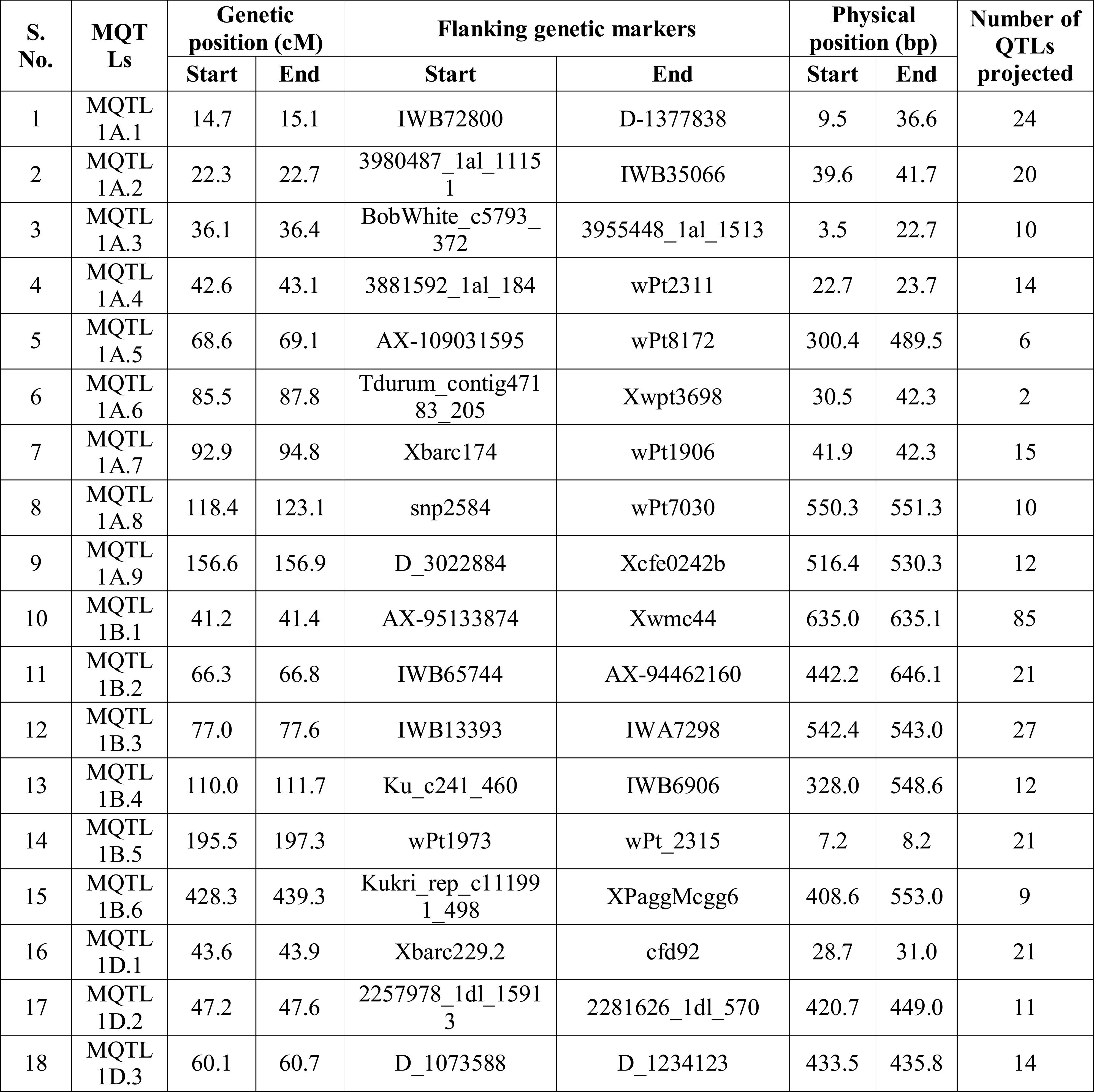
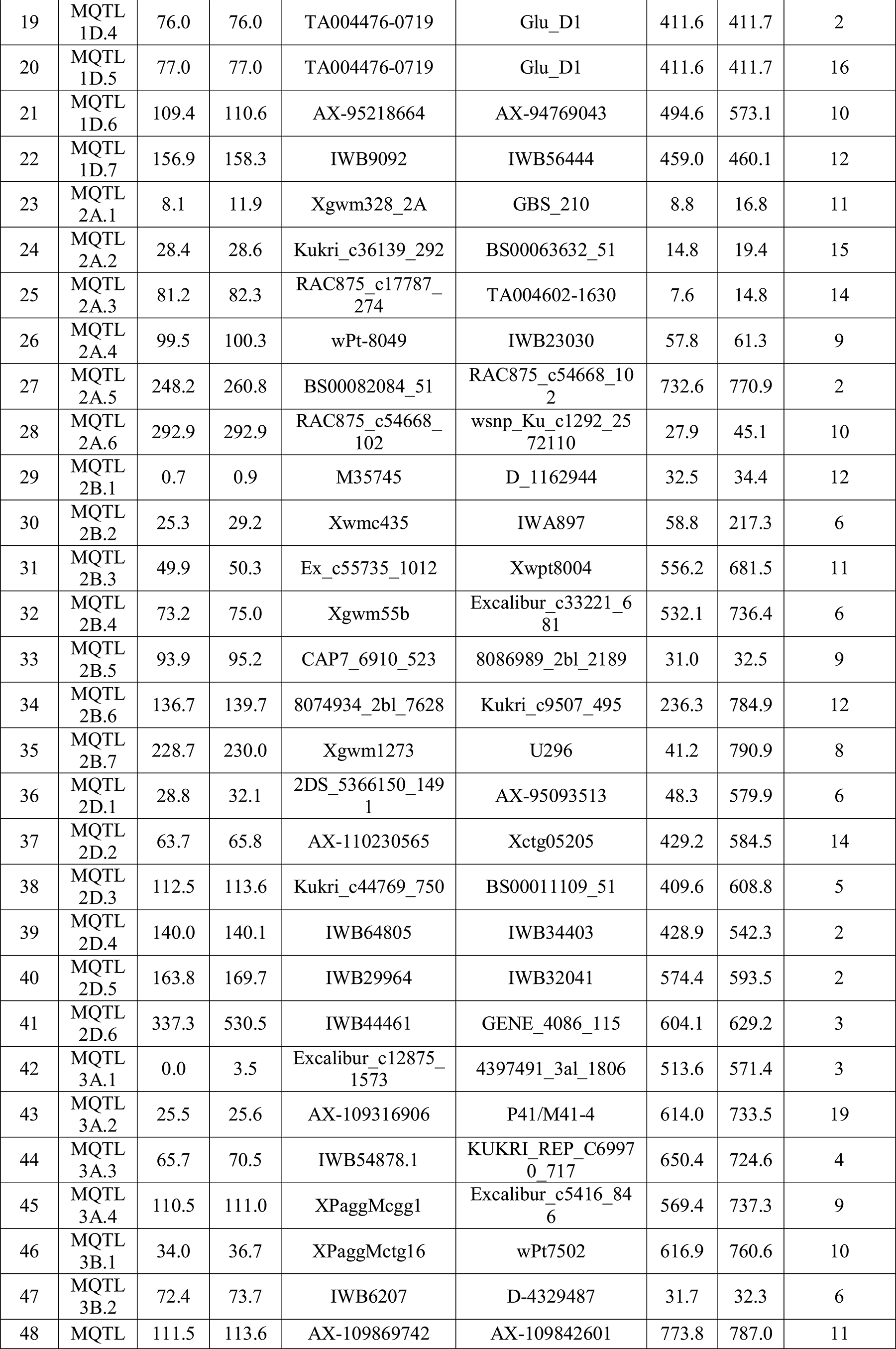
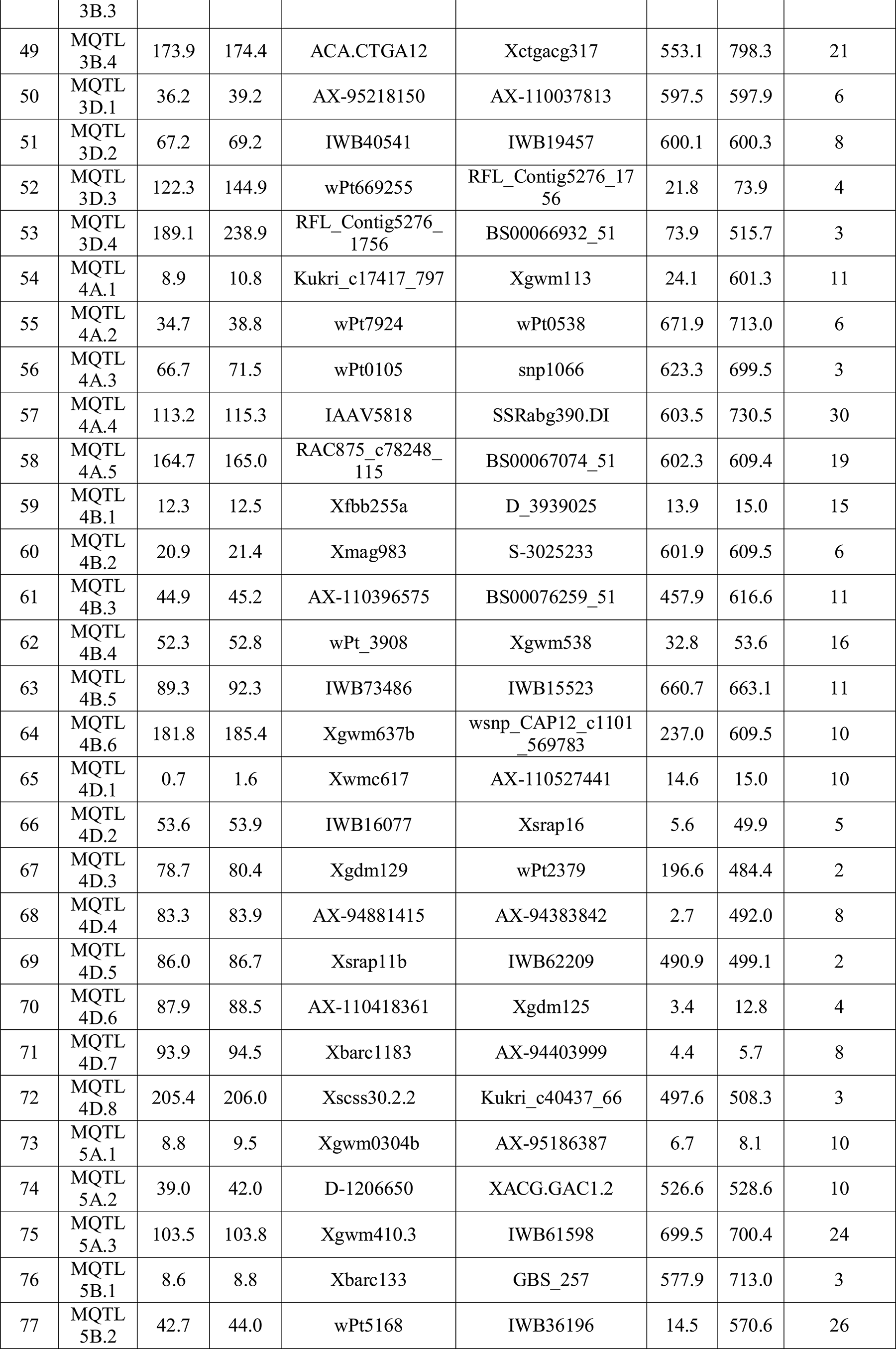
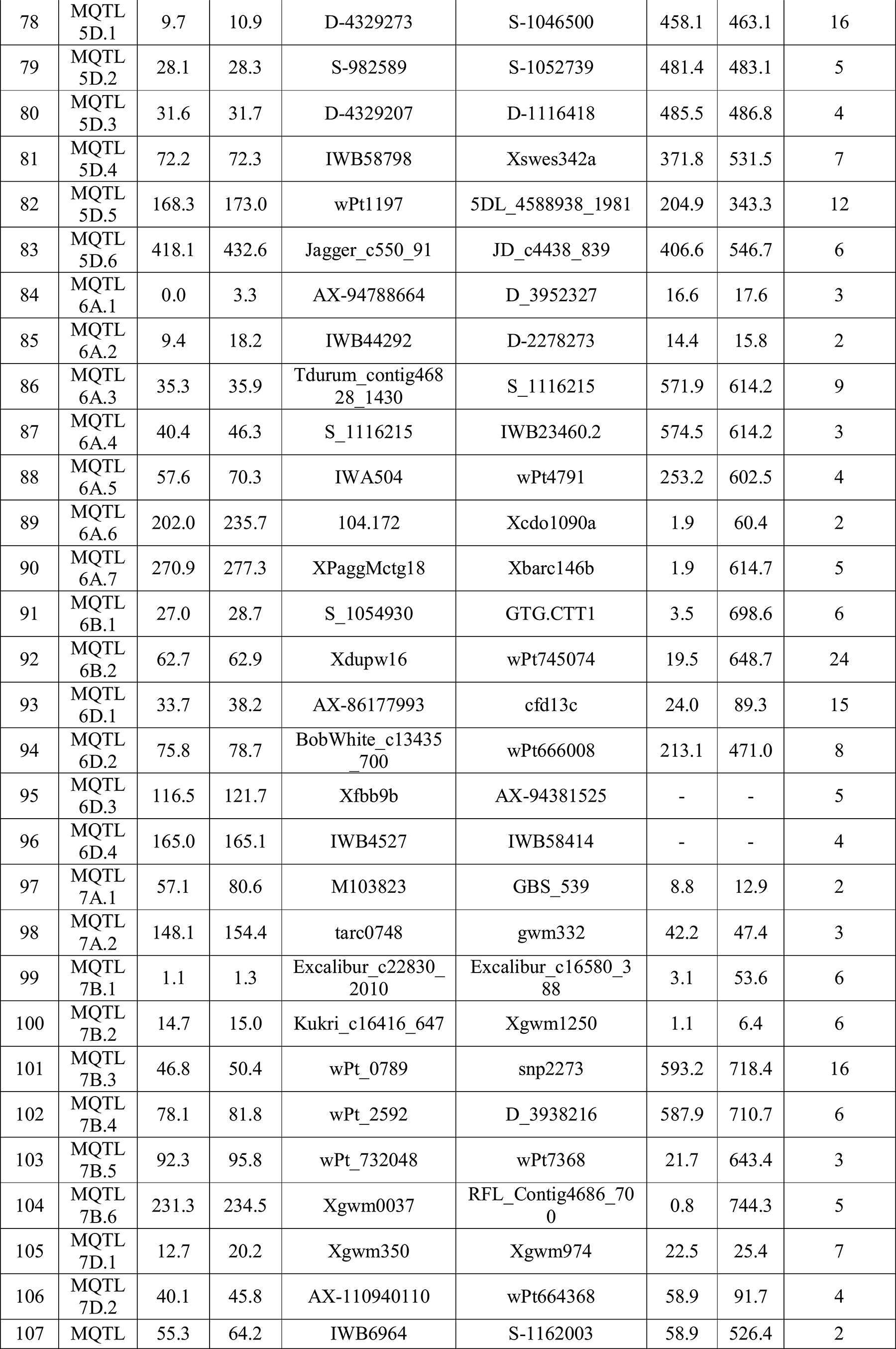
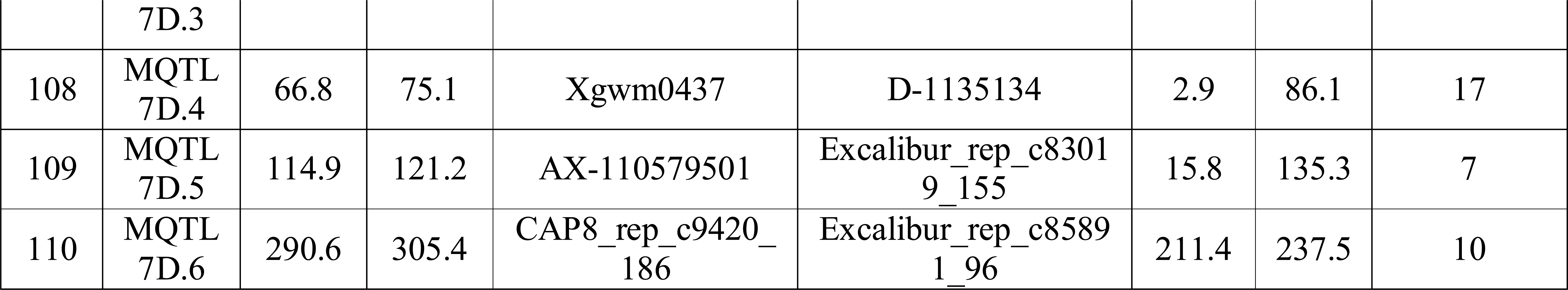
Summary of wheat MQTLs associated with different quality traits

Individual QTLs for processing quality traits were found to be associated with as many as 98 MQTLs and for that of the other traits are given in Fig 3b and S5 Table. All MQTLs were associated with at least one quality trait (Fig 3e). As many as 38 MQTLs were exclusively associated with only a single quality trait, whereas, four MQTLs (viz., 2D.2, 5A.2, 5D.4, and 7B.3) were found to be associated with 5 quality related traits (S5 Table). None of the MQTL included the QTLs associated with all of the six studied traits. MQTLs associated with multiple traits are considered to be more important for the breeding programmes aiming towards the simultaneous improvement of multiple quality traits. In this regard, MQTLs (viz., 2D.2, 5A.2, 5D.4, and 7B.3) were considered as more promising and breeders can use them to enhance multiple quality traits simultaneously.

### MQTLs validated with GWAS results

The physical coordinates of the MQTLs identified in the present study were also compared with MTAs reported in earlier GWA studies (S6 Table). Among the 108 MQTLs, as many as 43 MQTLs (39.81 %) co-localized with at least one MTA available from the GWAS (Fig 4; S6 Table). There were some MQTLs co-localized with MTAs available from more than one GWA study. For instance, MQTL6A 7 co-located with MTAs reported in seven different GWA studies. Each of the five MQTLs (viz., 1A.1, 1B.2, 3B.4, 5B.2 and 6B.2) involving at least 20 initial QTLs co-localized with multiple MTAs reported from different GWA studies.

Surprisingly, five genes, *TraesCS1A02G040600*, *TraesCS2D02G531100*, *TraesCS3B02G449200*, *TraesCS3D02G095700* and *TraesCS3D02G096000* described by Yang et al. (2021) GWA study were overlapped with four MQTLs identified in the present study. Among them, first three genes were present in the MQTL1A.1, MQTL2D.6, and MQTL3B.1, respectively. While, last two genes located on the 3D chromosomes were present in the MQTL3D.3. Functional annotation and GO study revealed their participation in the various biological processes associated with grain quality in wheat. Genes, namely *TraesCS3D02G095700* and *TraesCS3D02G096000* encode for the wheat allergens and trypsin and alpha-amylase inhibitor present in the seeds.

### Wheat homologues of known rice genes in MQTL regions

An extensive search made on known rice genes associated with quality traits led to the collection of information on 34 genes which were further utilized for the identification of their corresponding homologues in wheat MQTL regions. These genes encode proteins/products mainly belonging to the following families: starch synthase, glycosyl transferase, aldehyde dehydrogenase, SWEET sugar transporter, alpha amylase, glycoside hydrolase, glycogen debranching, protein kinase, peptidase, legumain, and seed storage proteins. Wheat homologues of 30 of these rice genes were identified in different MQTL regions (S7 Table); wheat homologues for three rice genes, namely, *RP6*, *RM1,* and *OsAGPL4/OsAPL4* could not be identified due to low sequence identity (<60%) and query coverage (<30%). Further, it has been noticed that, wheat homologues for nine rice genes are identified on the different chromosomes of wheat. For example, wheat homologues for rice gene, *Wx1* are present on the 4^th^ and 7^th^ chromosomes. Sixty-four wheat homologues of these 30 rice genes were detected in the 44 wheat MQTL regions (S7 Table); some MQTL regions harboured more than one wheat homologues of rice genes. For example, wheat homologues of rice genes *wx1*, *OsACS6* and *GBSSII* genes were present within the MQTL4A.2 (S7 Table).

### Ortho-MQTL analysis

Extensive investigation of orthologous gene models available from all the 108 wheat MQTLs (with known genomic coordinates) led to the identification of 23 wheat MQTLs (ortho-MQTLs) syntenic to 16 rice MQTLs (Fig 4; S8 Table). In some cases, more than one wheat MQTLs were found to be syntenic to a single rice MQTL. For instance, three wheat MQTLs (viz., 2B.3, 2D.2, and 2D.2) were found to be syntenic to a rice MQTL (i.e., MQTL4.5) (Figs 6 and 8). Conversely, single wheat MQTL was found to be syntenic to multiple rice MQTLs. For instance, wheat MQTL4A.4 was syntenic to two rice MQTLs located on chromosome 8 (viz., MQTL8.6 and MQTL8.8) and MQTL4B.6 was syntenic to two rice MQTLs located on chromosome 3 (MQTL3.3 and MQTL3.4).

Number of orthologous gene models in individual MQTL varied from 3 in MQTL1A.3 to 19 genes in MQTL2D.2. Five ortho-MQTLs (viz., 2A.1, 2D.2, 4D.8, 5D.1 and 7D.1) each involving more than 10 gene models syntenic to rice genome are presented in Fig 5. Gene models present in the two wheat MQTLs (viz., 2D.2 and 4D.8) located on chromosome 2D were found to be orthologous to rice MQTL4.5 and MQTL3.4, respectively, in the reverse order (Fig 5b, c). From this, it can be inferred that genomic regions in MQTL2D.2 and MQTL4D.8 were inverted once in the evolutionary time and conserved in the rice genome. MQTL5D.1 with 13 gene models was corresponding to rice MQTL3.7. Wheat genes present in MQTL5D.1 was found to be collinear to one rice MQTL3.7 (Fig 5d). We also reported a disruption in collinearity between the genes available from the wheat (MQTL7D.1) and rice (MQTL6.1) MQTLs (Fig 5e). Wheat MQTL2A.1 with 14 genes was found to be syntenic to the rice MQTL4.2 (Fig 5a).

**Fig 5.**
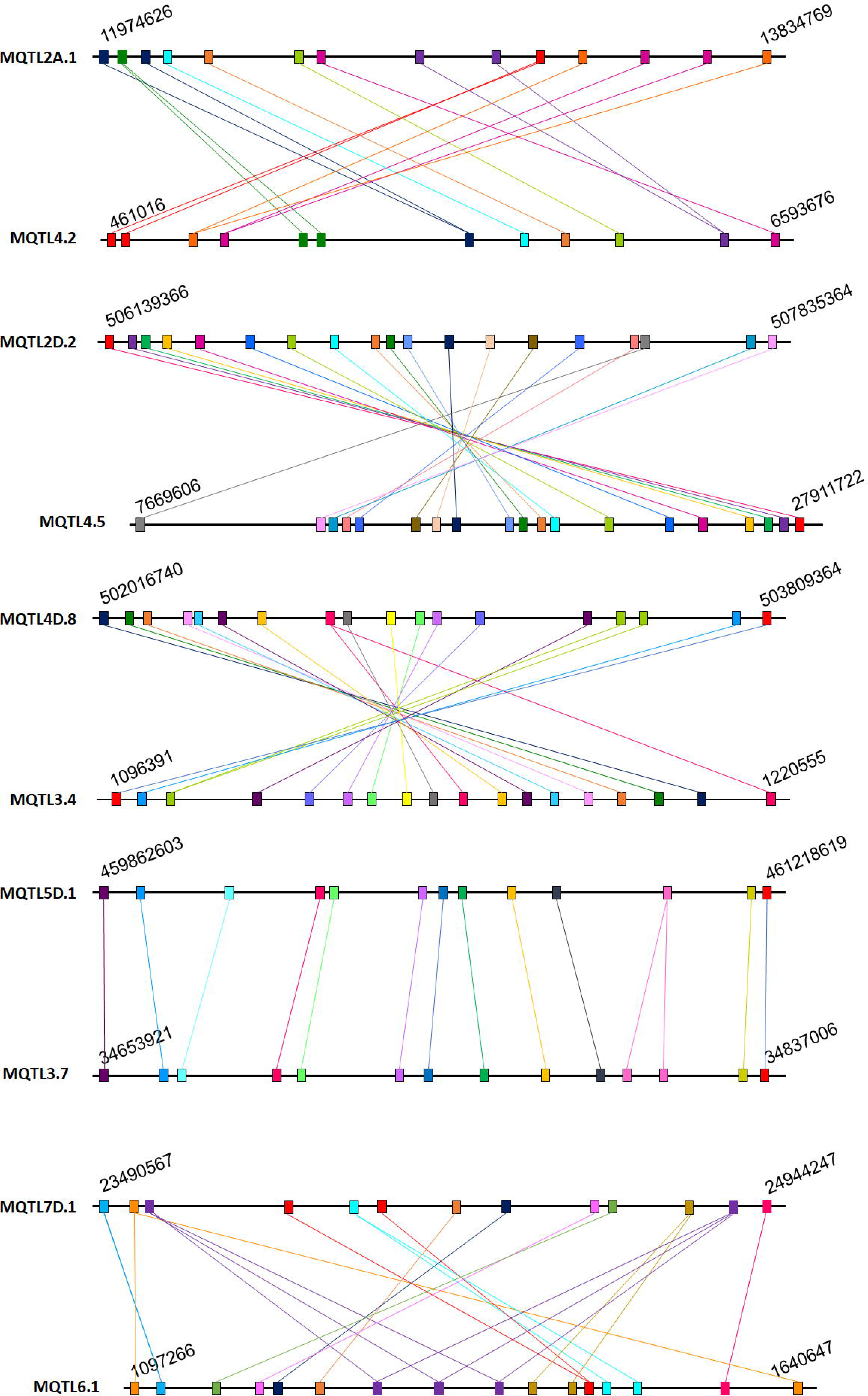
The syntenic region of MQTLs between the wheat and rice. The genomic position, chromosome number, and common genes between the wheat and rice are indicated.

### Gene models available in MQTL regions and their expression analysis

A total of 108 MQTLs were anchored to the physical map of wheat reference genome. Physical positions of the two MQTLs (viz., MQTL6D.3 and MQTL6D.4) could not be deduced, as these MQTLs were flanked by markers with no sequence information available. Gene mining within 108 MQTL regions permitted the identification of as many as 2533 gene models. Maximum number of gene models (276) were identified for MQTLs located on the chromosome 2A, while the minimum number of the gene models (6 gene models) were available from the MQTLs located on chromosome 6B (Table 2; S9 Table).

**Table 2.** Number of initial QTLs, projected QTLs, gene models identified in the different chromosomes

| **Chromosome** | **Initial number of QTLs** | **Projected QTLs** | **Unique gene models** | **Gene models with >2 TPM expression** |
| --- | --- | --- | --- | --- |
| 1A | 206 | 113 | 262 | 52 |
| 1B | 289 | 175 | 119 | 47 |
| 1D | 158 | 86 | 116 | 25 |
| 2A | 121 | 61 | 276 | 47 |
| 2B | 177 | 64 | 88 | 17 |
| 2D | 100 | 32 | 140 | 38 |
| 3A | 90 | 35 | 93 | 14 |
| 3B | 105 | 48 | 79 | 21 |
| 3D | 34 | 22 | 104 | 26 |
| 4A | 124 | 69 | 112 | 12 |
| 4B | 142 | 69 | 117 | 31 |
| 4D | 49 | 42 | 242 | 69 |
| 5A | 93 | 44 | 94 | 18 |
| 5B | 100 | 29 | 28 | 8 |
| 5D | 94 | 50 | 143 | 38 |
| 6A | 82 | 28 | 204 | 43 |
| 6B | 94 | 30 | 6 | 2 |
| 6D | 62 | 32 | 54 | 18 |
| 7A | 140 | 2 | 47 | 7 |
| 7B | 88 | 43 | 118 | 25 |
| 7D | 110 | 47 | 91 | 22 |
| **Total** | **2458** | **1121** | **2533** | **580** |

Expression analysis of available gene models detected 556 gene models with >2 transcripts per million (TPM) expression, which included 94 gene models with >5 TPM and 6 gene models with >10 TPM expression in grains and its component tissues such as seed coat, starchy endosperm, whole endosperm, aleurone layer and grain transfer cells in (Fig 8; S10 Table). Of these 556 gene models, 117 gene models showed constitutive expression in all the tissues studied. While, the remaining 439 gene models showed the differential expression (tissue specific). Further, 44 differentially expressed gene models with more than 5 TPM expression were involved in the improvement of grain quality (Fig 7). Based on their expression pattern, gene models could be divided into two classes, i.e., genes in class-I showed expression in the seed, while genes in the class-II exhibited expression in the different sub-tissues. Further, functional characterization of these genes revealed their involvement in transcriptional and translational regulation of various genes, signalling mechanism, metabolism, cellular development and transfer, etc.

**Fig 6.**
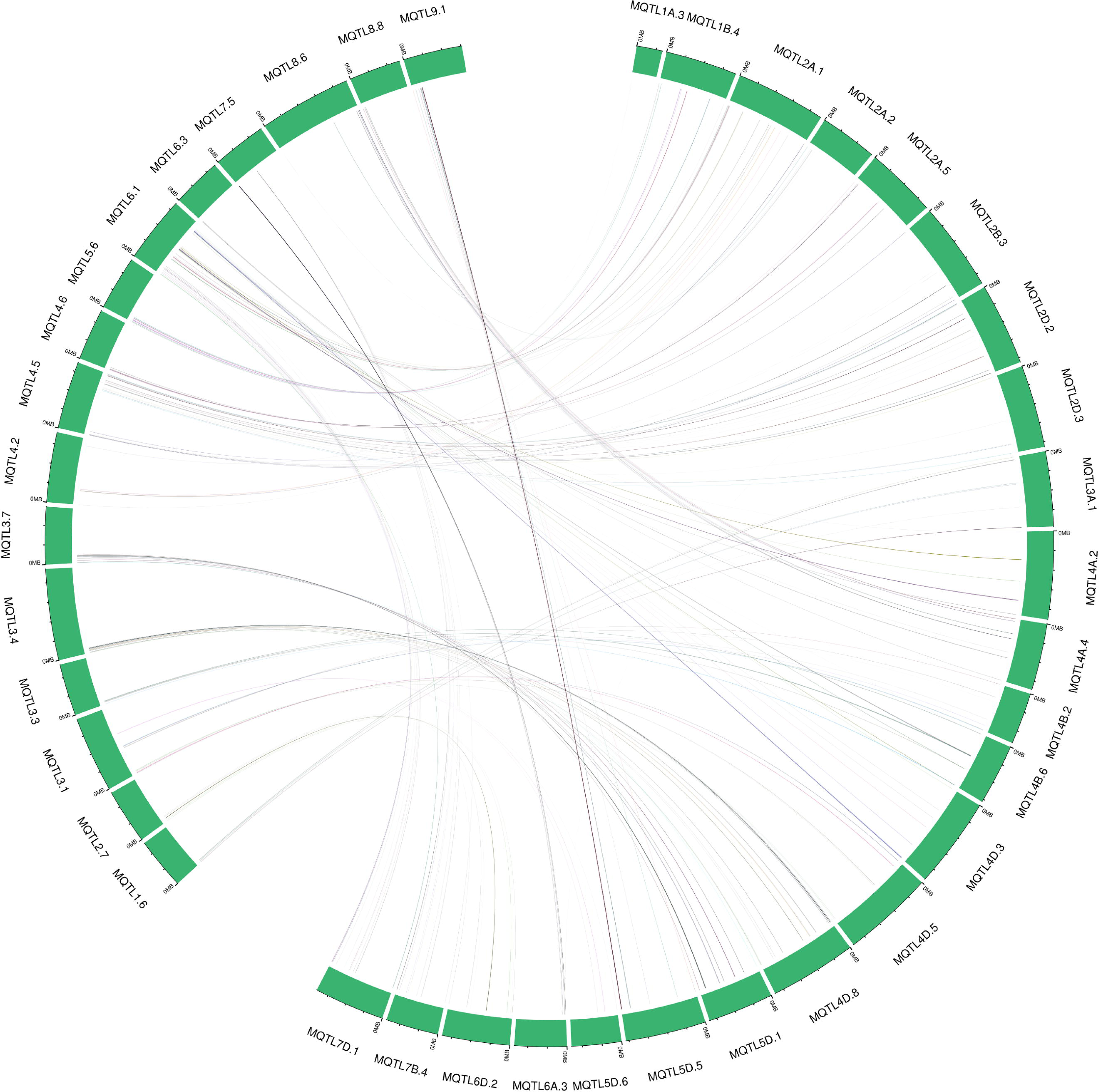
Circular diagram representing the conserved regions between wheat and rice MQTLs.

**Fig 7.**
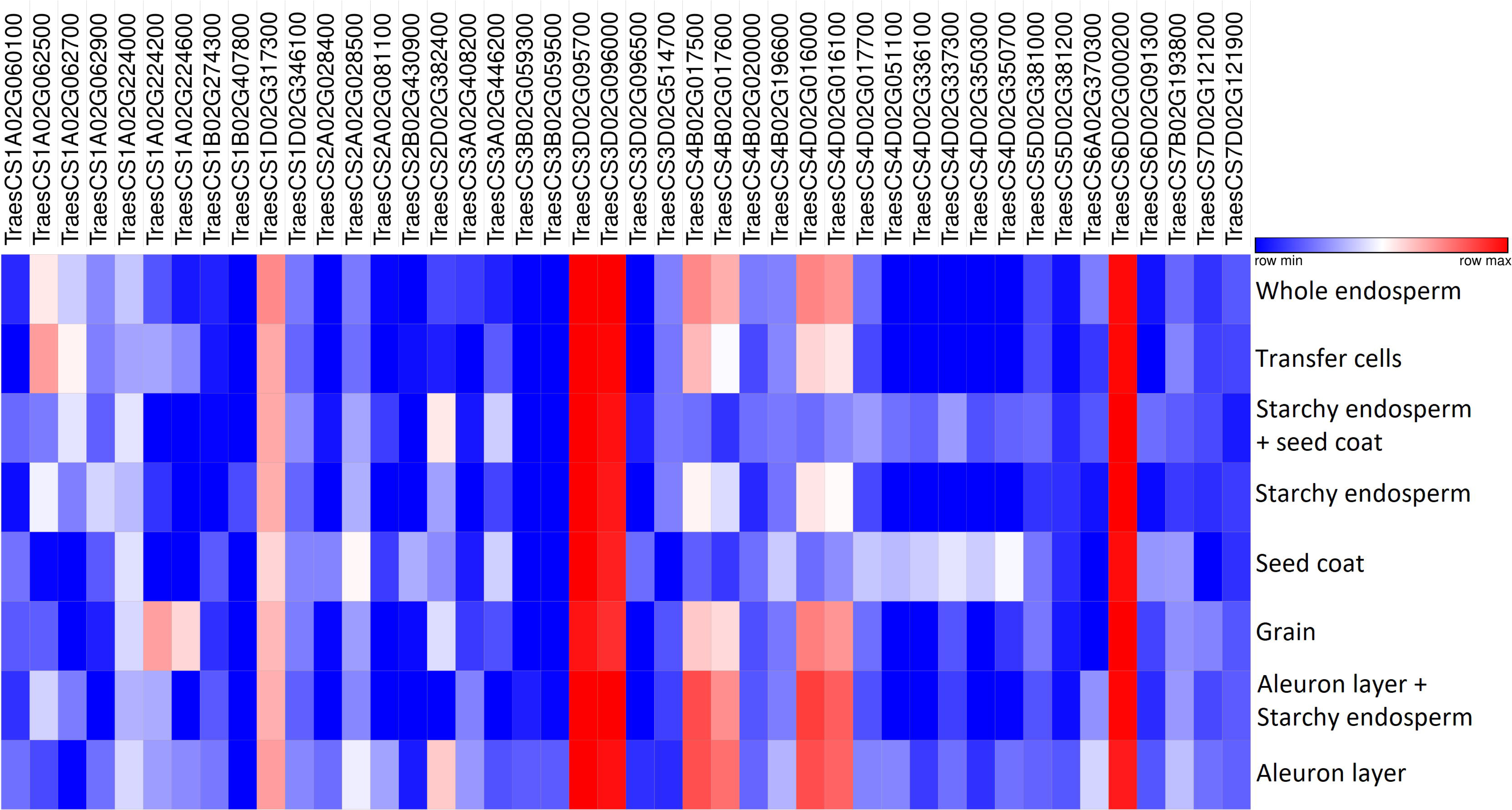
Expression pattern of 44 high-confidence candidate genes in the different relevant tissues.

**Fig 8.**
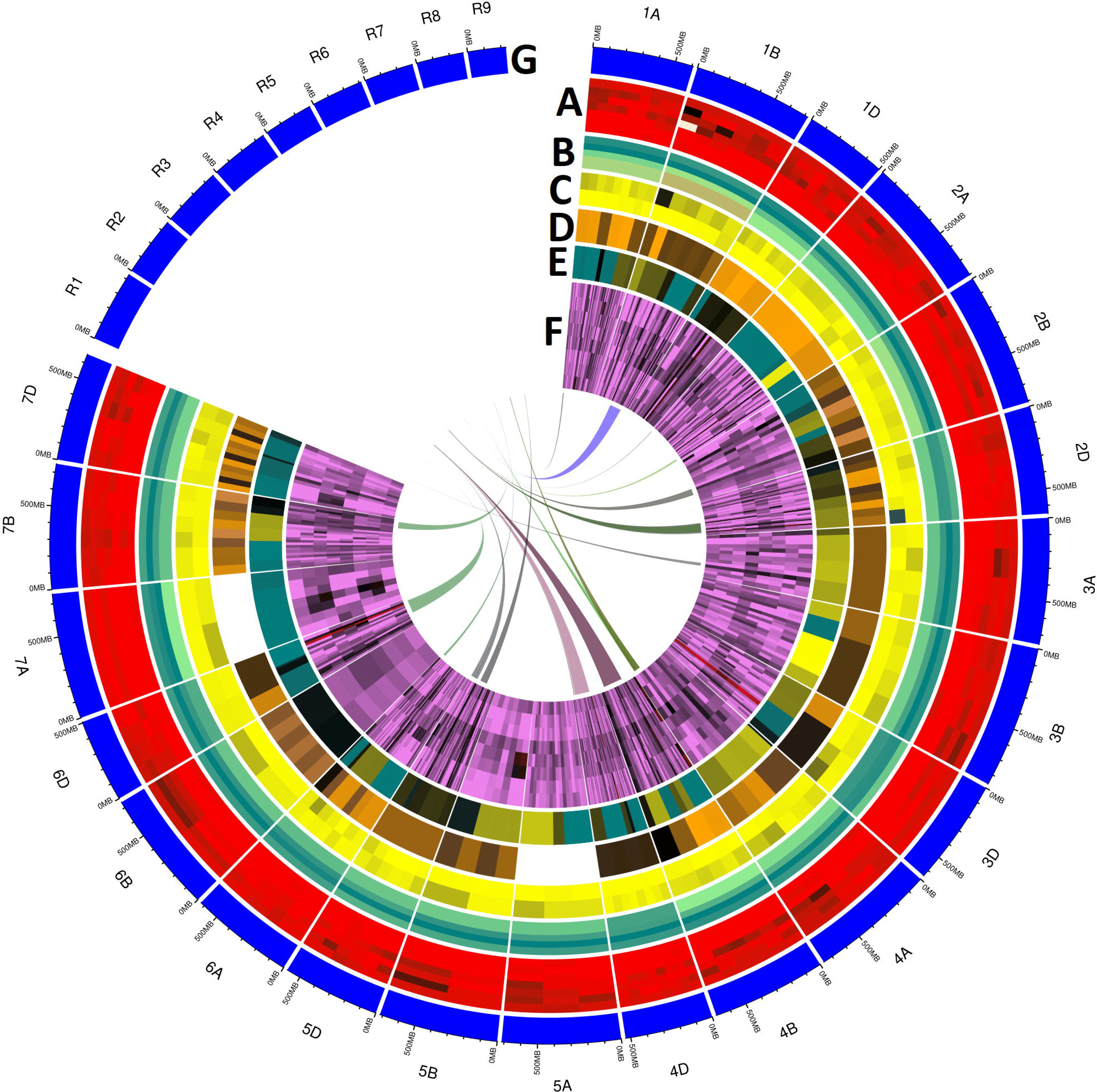
Circular diagram showing the different features of QTLs and MQTLs (A) number of QTLs for each trait in each MQTL (i.e., Ax, PPO, SV, DRP, nutritional traits, processing quality traits) and number of traits in each MQTL, **(B)** chromosome wise distribution of initial number of QTLs, projected QTLs, average confidence interval (CI) of MQTLs and average CI of initial QTLs, **(C)** number of initial QTLs projected on each of the MQTL, **(D)** MQTLs colocalized with known wheat genes, GWAS-MTAs and functionally characterized rice genes, **(E)** Gene density distribution in the MQTL regions, **(F)** expression pattern of candidate gene models with >2 TPM in the 8 different tissues (i.e., whole grain, whole endosperm, starchy endosperm, aleurone layer, seed coat, starchy endosperm + seed coat, transfer cells and aleurone layer + starchy endosperm), **(G)** syntenic region between wheat and rice MQTLs. Outer circle represents the physical maps of wheat and rice chromosomes (wheat chromosomes 1A to 7D and rice chromosomes R1 to R9).

In recent study, transcriptome analysis in the hexaploid wheat and its diploid progenitors identified the differentially expressed genes (DEGs) associated with carbohydrate metabolism (Kaushik et al. 2020). Seven DEGs (Table 3) identified from Kaushik et al. (2020) study were common to the genes characterized in the present study. Of these seven genes, six genes were downregulated (Table 3) and one gene, *TraesCS7B02G194000.1* was upregulated. Among the downregulated genes, two genes were encoding for NB-ARC domain containing proteins and one gene for the F-box domain (Table 3). While the functions of the three downregulated genes were not characterized.

**Table 3.**
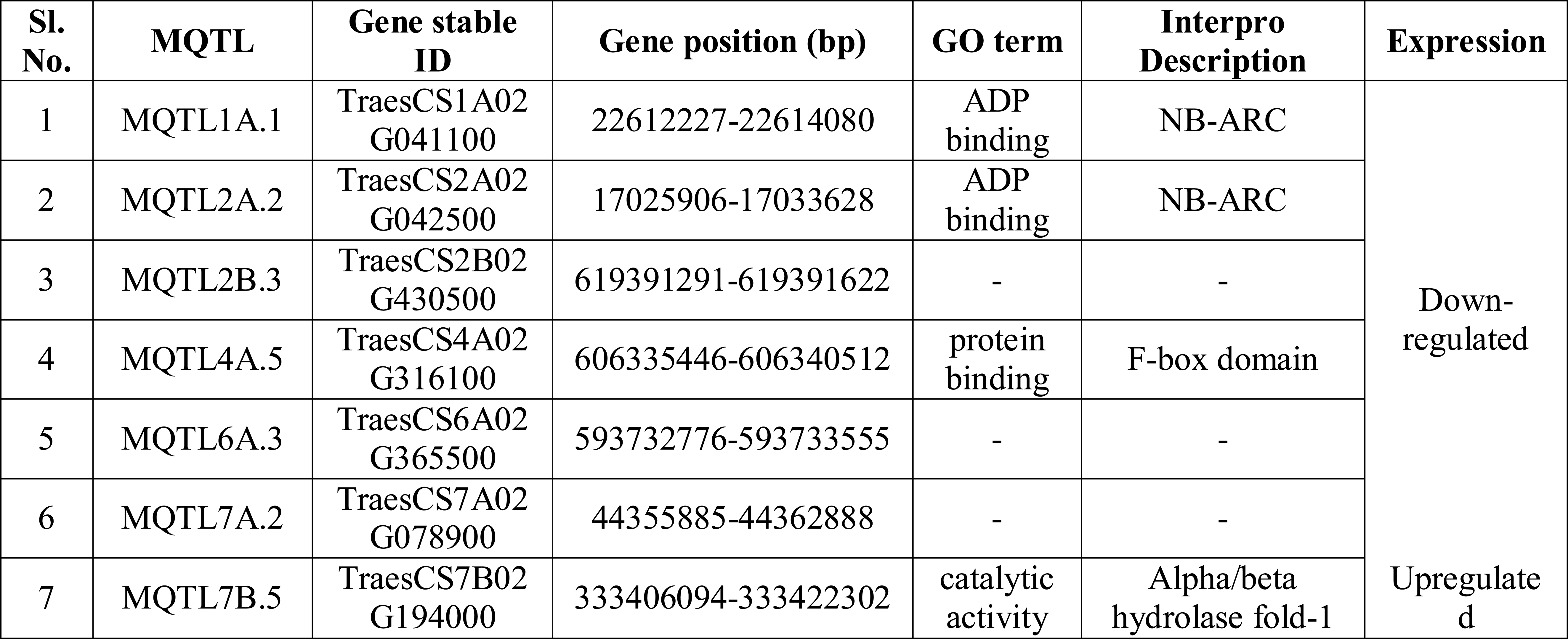
Differentially expressed genes detected from the transcriptomic studies (Kaushik et al. 2020) which co-localized with MQTLs identified in the current study

Cloned and characterized wheat genes associated with grain protein content (GPC) (*GPC-B1/NAM-B1*), HMW glutenin (*Glu-B1-1b*, *Glu-1D-1d*, *1Dx2t* and *Glu-1By9*) and starch synthase enzymes (*TaSSI*, *TaSSIIa*, *TaGBSSIa* and *TaSSIVb*) were identified in the 12 MQTL regions (S11 Table). Two MQTLs (6B.1 and 6B.2) present on chromosome 6B are associated with the GPC. MQTLs lying on the chromosome 1B (1B.2, 1B.3 and 1B.4) and 1D (1D.4 and 1D5) are act as a precursor for the HMW. Genes for the starch synthase enzyme present on the 1D (MQTL1D.2) and 7D (MQTL7D.2, MQTL7D.3, MQTL7D.4 & MQTL7D.5) chromosomes are responsible for the amylopectin biosynthesis via α also identified the 165 genes associated with catechol oxidase activity and metal ion binding (related PPO), Zn-transporter and zinc-binding site (Zn and Fe content), Small hydrophilic plant seed protein, Amino acid transporter and seed storage helical domain (seed storage protein), sweet-sugar transporters, UDP-glucuronosyl/UDP-glucosyltransferase and Sugar/inositol transporter (starch content) etc. Among them, 44 genes had the expression value of more than two TPM and were regarded as the putative CGs (Table 4). All the six CGs, *TraesCS3D02G095700*, *TraesCS3D02G096000*, *TraesCS4B02G017500*, *TraesCS4D02G016000*, *TraesCS4D02G016100* and *TraesCS6D02G000200* having high expression value of >10 TPM were showed to be involved in improving the seed quality through nutrient reservoir activity.

**Table 4.**
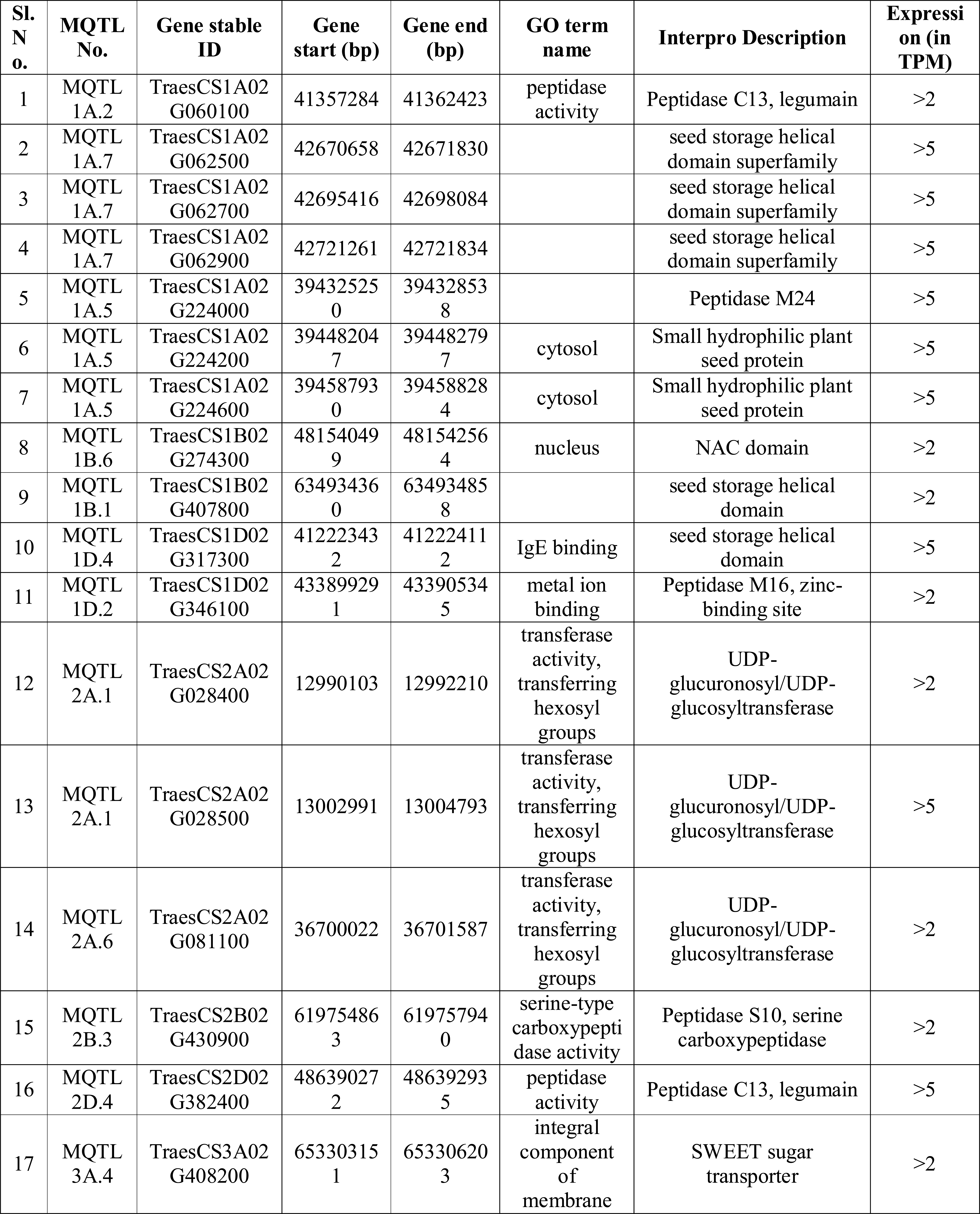
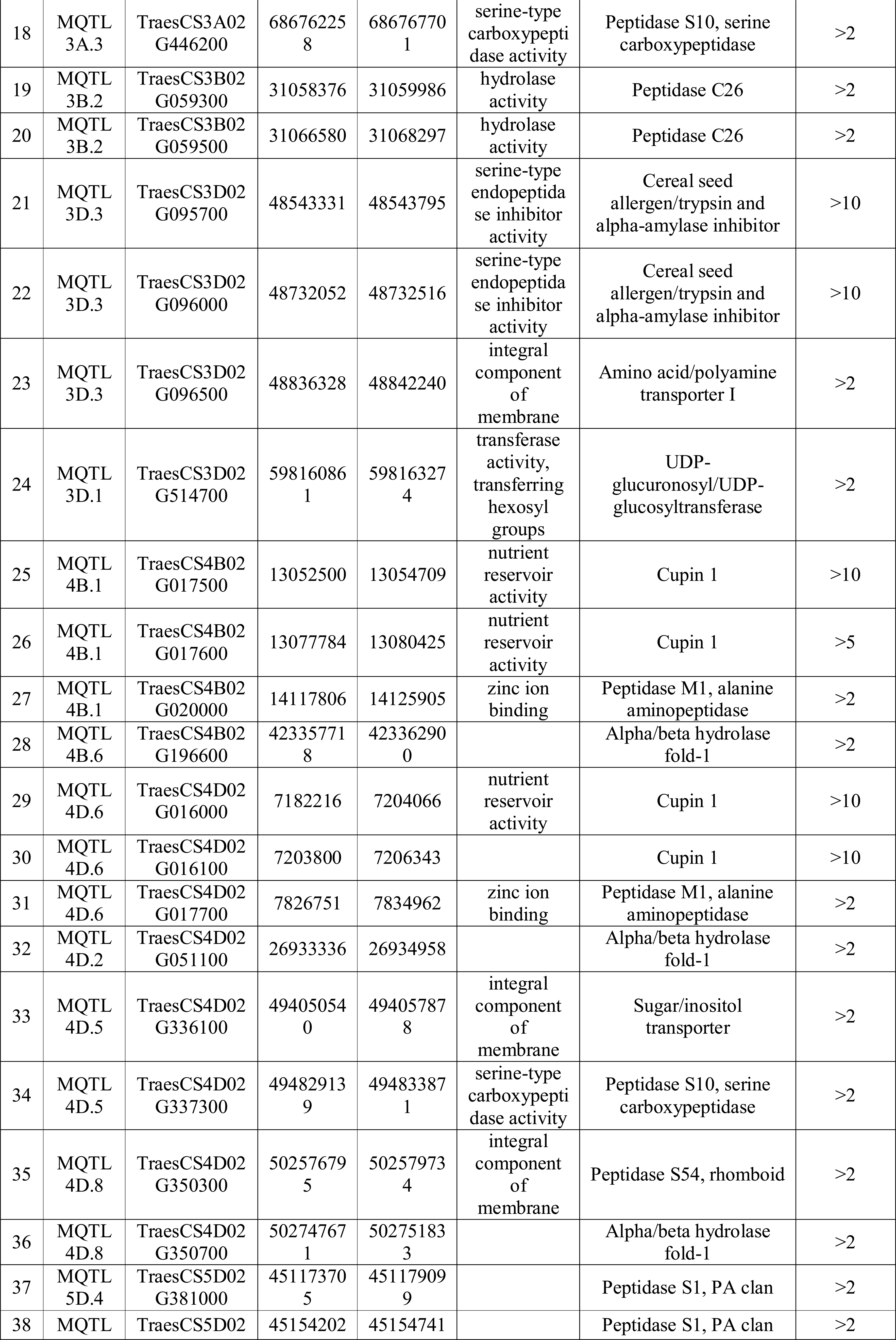
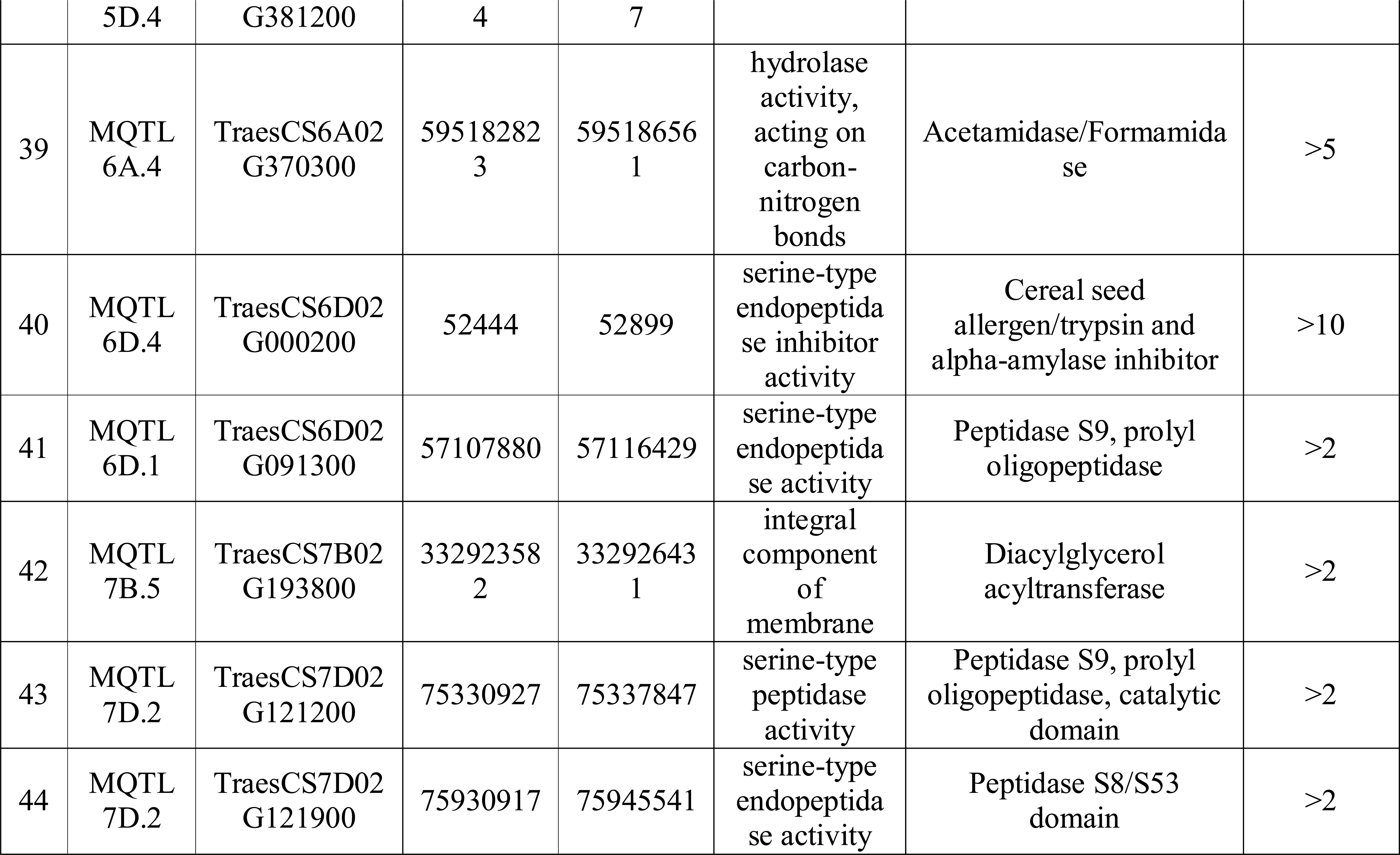
High-confidence candidate genes with >5 TPM expression identified in the present study

## Discussion

Discovery of molecular markers and advancements in QTL mapping strategies have promoted the plant researchers across the globe to underpin the complex genetic architecture of quality traits in wheat (Quraishi et al. 2017). Results of these intensive studies has accrued a large number of QTLs controlling the quality traits in wheat. Majority of these QTLs are minor QTLs (with low PVE value) and have larger CI, making them unsuitable for the marker assisted plant breeding (Kumar Saini et al. 2021). Further, it has been observed that, QTLs identified from one population may not be effective for a breeding programme involving the other population (Yang et al.). Downsides of these QTLs promoted us to reanalyze all of these loci together, with the goal of fine-tuning this vast amount of data, so that breeders and researchers can make greater use of these QTLs.

Meta-QTL analysis has the great capability to compile the information from multiple QTL mapping studies involving diverse environments with different genetic backgrounds to identify stable and reliable MQTLs with reduced CI (Welcker et al.). MQTL analysis have already conducted for the major crops including the rice, wheat, maize, etc. (Yang et al.; Li et al. 2013; Quraishi et al. 2017). Meta-QTL analysis for agronomic and quality traits in wheat has identified the six MQTLs for GPC and eight MQTLs for baking quality from the initial pool of 155 QTLs involving eight genetic studies published before 2013 (Quraishi et al. 2017). Further, seven MQTLs associated with Zn and Fe content were identified from initial set of 50 QTLs involving seven independent segregating populations. These studies have employed a small number of initial QTLs for the meta-analysis. However, the results of MQTL analysis have been found to be significantly and positively correlated with the quality of the results of meta-analysis (Yang et al.; Quraishi et al. 2017). Furthermore, as molecular genetics and QTL mapping methodologies are advancing, novel QTLs are adding to the database on a regular basis, therefore we must keep up with this pace in order to integrate new QTLs into more stable and reliable MQTLs.

Thus, to supplement the previous studies (Quraishi et al. 2017), in the present study, we conducted the meta-analysis for the QTLs reported during 2013-20 by projecting very large number of initial QTLs (1986 QTLs) on the high-density consensus map and identified 110 MQTLs each involving at least two initial QTLs. In addition to Zn, Fe, GPC and baking quality (studied by Quraishi et al. 2017), the present study also considered the other important quality parameters such as starch, PPO, Ax content, bread making properties, etc.

From the breeding perspectives, it is noteworthy to identify the most stable and reliable MQTLs involving a greater number of initial QTLs identified across the different populations and environments. Eleven such MQTLs involving more than 20 initial QTLs were detected in the present study. One MQTL located on chromosome 1B (MQTL1B.1) included as many as 85 initial QTLs, which is much higher than earlier study of (Quraishi et al. 2017). Further, to reduce the confidence interval, meta-QTL analysis relies on a large number of initial QTLs and heterogeneous populations (Berman and Parker 2002). Present study aggregated a vast QTL information from multiple genetic backgrounds and efficiently reduced the CI of the QTLs, thereby improved the accuracy of candidate gene prediction from the important QTL regions. The average CI of MQTL was 18.84-fold less than that of initial QTLs, and there were significant differences among different chromosomes. Some of the MQTLs with reduced CI identified in the present study may lay the foundation for molecular cloning and functional characterization of genes associated with quality traits in wheat. Further, this may offer the possibilities for the marker assisted gene transfer across the populations and contribute to the wheat quality improvement.

### Validating MQTLs with GWAS-MTAs

Genome wide association studies (GWAS) is a powerful tool which allows the investigation of highly complex characters by exploiting the recent and historical recombinant events present in the natural populations and permits the high-resolution mapping (Bush and Moore 2012). High-throughput, low-cost sequencing technologies have helped to identify MTAs for many quality traits in natural populations of wheat (Suliman et al.; Godoy et al. 2018). In the present study, 39.81% (43/108) of the identified MQTLs were verified by the GWAS-MTAs, which is in agreement with the earlier reports on the comparison of MQTLs with MTAs (Yang et al.; Aduragbemi and Soriano 2021). It is indicated from the previous studies (Yang et al.; Aduragbemi and Soriano 2021) and the present study, that, only a part of the MQTL region could be verified by the MTAs. Considering this, we propose the following two hypothesis: (i) none of the approaches whether meta-QTL analysis or GWAS approach included all the genetic variation present in the crop species, and (ii) genetic materials used in the two approaches were entirely different from each other. MQTLs could be considered more stable and consistent, if they are validated by more than one GWA study and also included the large number of initial QTLs from independent experiments. Five such MQTLs (1A.1, 1B.2, 3B.4, 5B.2 and 6B2) with more than 20 initial QTLs were verified with multiple MTAs derived from different GWA studies. These MQTL regions may be further investigated to explore the underlying complex genetics of the quality traits.

More interestingly, five genes identified in the wheat multi-locus GWA study (Yang et al.) were co-localized with four MQTLs detected in the current study. These genes are known to take active participation in the various biological processes to improve the grain quality in the wheat. For instance, gene, *TraesCS1A02G040600* encodes the cupin superfamily proteins such as phospho-glucose isomerase (PGI), which in turn takes part in the non-enzymatic protein storage in plant seeds (Dunwell and Gane 1998). Genes, *TraesCS3D02G095700* and *TraesCS3D02G096000* encode for the wheat allergens and trypsin & alpha-amylase inhibitor, respectively. Wheat allergens such as seed storage proteins (glutelin and prolamins) causing celiac disease may reduce the nutritional value of wheat seeds. Creating the null mutants for such genes through RNAi or CRISPR-CAS9 may reduce such allergic compounds present in the seeds (Zhang et al. 2014). Alpha-amylase inhibitors prevent the hydrolysis of storage starch granules present in the endosperm and there by maintain the integrity of starch in the seeds (Ali and Elozeiri 2017).

### Ortho-MQTL mining: revealing conserved genomic regions between wheat and rice

Identification of ortho-MQTLs among the related crop species may provide a better understanding of genes controlling various quality characters with similar evolutionary lineage and conserved functions. Further, this may provide potential opportunities to transfer information from on species to another (Khahani et al. 2020; Kumar Saini et al. 2021). Ortho-MQTL analysis have been sparingly conducted in wheat, with only three studies are available for different studies including, nitrogen-use efficiency and related traits (Quraishi et al. 2017; Kumar Saini et al. 2021) and thermotolerance (Kumar et al. 2020). However, to our knowledge, no study has reported the ortho-MQTLs for the quality related traits in wheat and other cereals so far. This is the first report, where we identified the 23 ortho-MQTLs associated with quality traits conserved between wheat and rice. Functional characterization of gene models present in the wheat ortho-MQTLs revealed their associations with the sugar-phosphate transportation, seed storage protein, lipid storage activities, etc. Wheat gene, *TraesCS1B02G248000* present on the ortho-MQTL1B.4 is belonging to the NAC domain family and is associated with improving the GPC in wheat seeds. Corresponding orthologous rice gene, *BGIOSGA019745* located on MQTL5.6 belongs to the same gene family and is associated with improving the seed size (Mathew et al. 2016). Further, *TraesCS2B02G430300* gene present on the ortho-MQTL2B.3 is belonging to hydrophobic seed protein domain which accumulates hydrophobic proteins (HPS) on the seed surface. HPS are the abundant seed constituent and a potentially hazardous allergen that causes asthma in persons allergic to the crop dust (Gijzen et al. 1999). Phenotypic or genetic screening may be devised to select plants with reduced amounts of HPS on the seed surface and may offers new avenue for lessening the health hazard of seed dust exposure. Orthologue for this gene, *BGIOSGA016898* is present on the rice MQTL4.5 and is belong to the same class of protein domain. Exploring of such conserved genomic regions among related species may unravel the common regulatory pathways and may assist the utilization of genomic resources from the model species, which may greatly contribute to the rapid crop improvement.

### Candidate genes available in MQTL regions

MQTLs are considered as the potential targets for mining candidate genes (CGs) associated with the traits in question. Further, MQTL regions have been shown to have high correlation with gene density in genome as revealed by earlier reports on meta-QTL studies conducted in different crops including rice, wheat, and maize (Yang et al.; Swamy et al. 2011; Quraishi et al. 2017). In wheat, previous studies have reported 15,772 gene models for yield, baking quality and grain protein content (Quraishi et al. 2017), 324 gene models for Fusarium head blight (Venske et al. 2019), 228 gene models for drought (Kumar et al. 2020), and 237 gene models (Yang et al.) for yield related traits. In the present study, gene mining within 108 MQTLs identified 2533 gene models in wheat, at least some of them are known to be associated with quality traits.

Gene expression analysis is the study of how genes are transcribed to produce functional gene products, such as RNA or protein. *In silico* expression analysis permitted the identification of 556 gene models with more than 2 TPM expression. An extensive survey of literature revealed their involvement in transcriptional and translation regulation of different genes, signalling mechanism, metabolism, cellular development and transfer etc. (Table. S7). In the present study, ABC transporter genes were identified on chromosomes 1A, 2A, 2B, 4A, 4B, 6D and 7B, which may give opportunity to develop nutritionally enriched, high-yielding *wheat* cultivars. ABC type transporters are known to regulate seed weight, by transcriptional regulation and modulation of the transport of glutathione conjugates in chickpea (Basu et al. 2019). Along with seed weight, they may enhance the quality component traits, as reported in chickpea (Basu et al. 2019). Ammonium transporter genes available in MQTL6A.5 may play a crucial role in the transport of seed nitrogen (N) and metabolic processes for seed N accumulation, seed yield and N use efficiency (NUE) (McDonald and Ward 2016). They take part in increasing the level of specific amino acids, total soluble protein, and specific high-quality proteins in the seeds. Overall, the present study identified at least 165 gene models associated with catechol oxidase activity and metal ion binding (related PPO), Zn-transporter and zinc-binding site (Zn and Fe content), small hydrophilic plant seed protein, amino acid transporter and seed storage helical domain (seed storage protein), sweet-sugar transporters, UDP-glucuronosyl/UDP-glucosyltransferase, sugar/inositol transporter (starch content) etc. which are known to be associated with quality related traits in different crops. 44 of these 165 gene models with functions previously reported to be important for quality related traits in different crops were selected and recommended for future basic studies involving cloning and functional characterization.

Gene expression analysis of hexaploid bread wheat through RNA-seq and the GeneChip®Wheat Genome array identified numerous DEGs related to starch biosynthesis, nutrient reservoir activity, carbohydrate metabolism and seed storage protein synthesis (Kaushik et al. 2020). Seven DEGs identified in (Kaushik et al. 2020) study were co-located with some MQTLs identified on following 7 chromosomes 1A, 2A, 2B, 4A, 6A, 7A and 7B in the present study. Of these seven genes, six genes were downregulated and one gene, *TraesCS7B02G194000.1* was upregulated. Among the downregulated genes, two genes encode for NB-ARC domain containing proteins and one gene for the F-box domain (Table 3). While the functions of the three downregulated genes have not been characterized. Up-regulated gene, *TraesCS7B02G194000* encode the alpha/beta hydrolase enzyme which take part in the multitude of biochemical processes, including bioluminescence, fatty acid and polyketide biosynthesis and metabolism (M 2000).

To strengthen the location of MQTLs identified in this study, a search for co-localization of quality related genes was also carried out. This enabled detection of functionally characterized wheat genes such as, *GPC-B1/NAM-B1*, *Glu-B1-1b*, *Glu-1D-1d*, *1Dx2t*, *Glu-1By9*, *TaSSI*, *TaSSIIa*, *TaGBSSIa* and *TaSSIVb* in different MQTL regions. MQTL6B.1 and MQTL6B.2 were identified to be associated with the GPC, which may also take part in the nutrient remobilization from leaves to developing grains (Distelfeld et al. 2014). Genomic regions underlying the five wheat MQTLs, (viz., 1B.2, 1B.3, 1B.4, 1D.4 and 1D.5) carried some known genes which are the known precursors of HMW glutenin, a prime deciding factor for determining the dough elasticity and bread-making quality in wheat (Anjum et al. 2007). Genes for the starch synthase enzyme available from MQTL1D.2, 7D.2, 7D.3, 7D.4 and 7D.5 regions are responsible for the amylopectin biosynthesis via α glycosidic linkages. Amylopectin-A is the unique chemical compounds present in the wheat which trigger the low-density lipoproteins, which take part in the transportation and delivery of fatty acids, triacylglycerol, and cholesterol in many plant organs (Horstmann et al. 2017).

### Wheat homologues of known rice genes available in MQTL regions

Comparative genomics study of wheat with model grasses such as rice has revolutionized the molecular genetics and contributed to the wheat improvement by identifying the linkage blocks, gene rearrangements and conserved regions in the wheat (Kumar et al. 2020). Functionally characterized rice genes are known to have similar function in the wheat and their homologues have been identified to be co-located with MQTLs for the various traits (Hanif et al. 2016). For instance, in the present study, three wheat homologues of one rice genes (*OsSWEET4*) were identified in different MQTLs located on homoeologous group 2 chromosomes, these wheat homologues may be engaged in the embryo nourishment by supplying the sufficient amount of nutrients to the developing embryo in the wheat (Yang et al. 2018).

Using comparative genomics, homologues of these genes in wheat can be characterized, and functional markers for these genes can be developed and validated. A meta-analysis of QTLs associated with grain weight in tetraploid wheat, for instance, resulted in the identification of one important locus, mQTL-GW-6A on chromosome 6A (Avni et al. 2018). Within this MQTL region, authors discovered and characterized a wheat homologue of the rice gene, *OsGRF4* (Avni et al. 2018). This suggests that combining a MQTL study with a well-annotated genome can result in the rapid identification of CGs underlying traits of interest. Manipulation and integration of these genes in the breeding programme may contribute to the enhanced wheat quality.

## Conclusion

Meta-analysis of QTLs and comparative genomic approaches helped us to dissect the complex genetic architecture underlying various quality parameters such as GPC, total starch, Zn, Fe, PPO, baking and bread making properties in the wheat. Identified MQTLs and corresponding high-confidence CGs may enhance the marker assisted breeding for quality improvement in wheat. Molecular cloning and functional characterization of these CGs may further improve the understanding of wheat genetics. Ortho-MQTLs identified in this study may help to understand the common evolutionary pathways underlying the quality traits. Breeders across the globe can make use of most promising MQTLs (viz., 2D.2, 5A.2, 5D.4, and 7B.3) associated with multiple quality traits and CGs in their study to improve the quality related traits in wheat.

## Supporting information

S1 Table. List of identified QTLs from the 50 (63 populations) independent studies and the related information (QTLs highleted with yellow colour have not been used for MQTL analysis)

S2 Table. Summary of the individual QTL mapping studies utilized in the present study S3 Table. Traits grouped into different major trait categories

S4 Table. Wheat_Reference_GeneticMap-2021

S5 Table. Details on identified MQTLs detected in the present study

S6 Table. MQTLs co-located with MTAs identified in the earlier GWA studies

S7 Table. Wheat homologues of known rice genes identified in different MQTLs located on sub-genomes A, B and D

S8 Table. Ortho-MQTLs identified between wheat from rice

S9 Table. Details on gene models identified in the present study

S10 Table. List of important gene models and their expression pattern in different tissues (MQTLs in black had >2 TPM, MQTLs in red had >5 TPM, and MQTLs in blue had >10 TPM expressions

S11 Table. Functionally characterized wheat genes co-localized with MQTLs identified in the current study

## Declarations

### Availability of data and material

Relevant data are included in this paper and its associated Supplementary Information (SI).

### Code availability

Not applicable

### Author’s contribution (SG, DKS, GS, PH, PK, MS, MJT, AS)

AS, SG, and DKS conceived and planned this study. SG, PH, PK, and MJT performed the literature search, retrieved data, developed consensus map, conducted meta-analysis. GS, DKS and MS helped SG in the analysis and interpretation of results and in writing of the first draft of the manuscript. AS, DKS and GS critically revised and edited the manuscript. All authors have read and agree to the final version of the manuscript.

### Ethics approval and consent to participate

Not applicable.

### Consent to participate

Not applicable.

### Consent for publication

Not applicable

